# Predicting Cell-Penetrating Peptides: Building and Interpreting Random Forest based prediction Models

**DOI:** 10.1101/2020.10.15.341149

**Authors:** Shilpa Yadahalli, Chandra S. Verma

**Affiliations:** Bioinformatics Institute A*-STAR (Agency for Science, Technology and Research), Singapore; Department of Biological Sciences, National University of Singapore, Singapore; School of Biological Sciences, Nanyang Technological University, Singapore

## Abstract

Targeting intracellular pathways with peptide drugs is becoming increasingly desirable but often limited in application due to their poor cell permeability. Understanding cellular permeability of peptides remains a major challenge with very little structure-activity relationship known. Fortunately, there exist a class of peptides called Cell-Penetrating Peptides (CPPs), which have the ability to cross cell membranes and are also capable of delivering biologically active cargo into cells. Discovering patterns that make peptides cell-permeable have a variety of applications in drug delivery. In the current study, we build prediction models for CPPs exploring features covering a range of properties based on amino acid sequences, using Random forest classifiers which are often more interpretable than other ensemble machine learning algorithms. While obtaining prediction accuracies of ~96%, we also interpret our prediction models using TreeInterpreter, LIME and SHAP to decipher the contributions of important features and optimal feature space for CPP class. We propose that our work might offer an intuitive guide for incorporating features that impart cell-penetrability into the design of novel CPPs.

## 1 Introduction

Cell-penetrating peptides (CPPs) are short peptides normally observed to be around 5-30 amino acids in length, which have an ability to enter the cells without irreversibly damaging the cell membrane (Milletti, 2012). They can also carry cargoes ranging from probes or therapeutics which can be small molecules, peptides or proteins into cells (Li et al., 2015). These properties make them attractive as potential drug delivery agents (Milletti, 2012; Heitz et al., 2009). While this method of cellular delivery is desirable, it is also associated with issues such as toxicity and immunogenic responses that render them undesirable (Dinca et al., 2016). Majority of therapeutically interesting peptides fail to permeate the cells, unable to find their protein targets. Peptide as drugs are more desirable as compared to small molecules especially when the target is a protein-protein interaction site and peptides exhibit higher specificity. Hence there is a need to understand the relationships between the sequences of the peptides and their ability to penetrate the cells.

Over the years, with the increase in the availability of biological data for CPPs, several machine learning (ML) based predictors have emerged which are summarized elsewhere (Wei *et al.*, 2018; Hansen *et al.*, 2008; Su *et al.*, 2019). Mining the characteristics of peptides such as amino acid composition, biochemical properties and many novel feature representation methods have been used in several predictors to obtain accuracies higher than 80%. While much progress has been made in developing new prediction algorithms, only a few studies have focused on understanding the feature contributions and optimal feature space of CPPs. This can help us create strategies for designing CPPs and introducing cell-penetrability into other peptides of our interest. Towards this aim, we review the previously available datasets to construct robust training data, build random forest-based prediction models and interpret the models using various methods of model explainability like TreeInterpreter, LIME and SHAP in an attempt to understand optimal feature space in CPPs. Interpreting prediction models is also necessary to build more trust in them. We test the sensitivity of our models to various feature vectors, class imbalance and sequence similarities in the training data. We also discuss a prediction model built for non-cationic CPPs and analyze feature preference in them as compared to the cationic class of CPPs. In the end, we discuss further challenges in the prediction of CPPs along with a few discrepancies which we found in the current datasets which might arrive due to different experimental conditions.

We present a faster, simpler and interpretable prediction model, without compromising on the accuracy and we believe this work helps in increasing our understanding of CPPs.

## 2 Methods

### 2.1 Algorithm Selection

We have built ML-based classification models for the prediction of CPPs. We chose a random forest classifier (RFC) (Breiman, 2001) which has been proven to be effective for classification tasks in many fields of computational biology (Wei, Xing, *et al.*, 2017; Chen *et al.*, 2015). It is quite robust in handling various data types at different scales and is resistant to overfitting (Trevor Hastie, Robert Tibshirani, 2009; Bénard *et al.*, 2019). The main reason for us to choose RFC however is that they are often more interpretable and easier to analyze than many other ensemble ML algorithms (Ishwaran, 2007; Louppe, 2014). The RFC algorithm comprises of an ensemble of decision trees, each of which is grown by a subset of features selected from the input feature vector. The number of features for each tree is determined by multiple factors, such as the generalization error, classifier strength, and inter-dependence within them (Breiman, 2001). We used scikit-learn’s RFC library (Pedregosa *et al.*, 2011) to build our models. The parameters used to build RFC models are obtained by hyperparameter tuning method, GridSearchCV, from scikit-learn. These parameters are mentioned in Supplementary information section 1 (SI-S1).

### 2.2 Dataset construction

We have used the following datasets in our study (these are also listed in Table 2).

- Dataset A. Dataset from CellPPD (Gautam *et al.*, 2013), downloaded from CPPsite 2.0 (Agrawal *et al.*, 2016). There are a total of 1416 sequences in this dataset with an equal number of CPPs and non-CPPs.
- Dataset B. ‘*Benchmark*’ dataset from CellPPD. These sequences referred by authors (Gautam *et al.*, 2013) as Benchmark dataset has a total of 343 sequences with 136 CPPs and 207 non-CPPs.
- Dataset C. Dataset from SkipCPP-Pred (Wei, Tang, *et al.*, 2017) has a total of 924 sequences with an equal number of CPPs and non-CPPs.
- Dataset D. Dataset from KELM-CPPpred (Pandey *et al.*, 2018) has a total of 826 sequences with an equal number of CPPs and non-CPPs.

We used these datasets as they are from state-of-the-art predictors, have high levels of predictive performance and the datasets are publicly available.

Dataset E. Ensemble dataset: We build this dataset by combining above 4 datasets. Most of the CPPs from the above datasets are from CPPsite 2.0 database, which is a golden source for CPPs, hence we remove duplicate sequences after combining the above datasets. Further, we remove a few entries with discrepancies in their labels across the sources (these entries are listed in SI-S2) and we are left with 955 CPPs. This dataset is available on request.

Since it has been well established that balanced datasets perform better and that imbalanced datasets present several different problems in ML methods (Sanders *et al.*, 2011), we try to maintain a similar number of CPP and non-CPP entries while training. However, we have also compared the model performances with/without rebalancing the data points.

We also test the sensitivity of the model to sequence similarities in the training data by building prediction models with/without sequence redundancy removal at various sequence identity cutoffs. It should also be noted that we do not have sequences containing modified or non-natural amino acids in our datasets.

### 2.3 Feature engineering

Following types of feature vectors are used as an input to our RFC:

1. Amino Acid Frequencies (AAF)
2. DiPeptide Frequencies (DPF)
3. TriPeptide Frequencies (TPF)
4. BioChemical Properties (BCP)
5. Ensemble-feature vector (EnF): by combining top-scoring features from these four models. We consider AAF, as it is known that certain types of residues are found with a higher frequency in CPPs, as outlined in the compositional based model (Garg *et al.*, 2005). DPF and TPF encapsulate information related to neighbouring residues, thus bringing in effects due to the order of amino acids (Petrilli, 1993). All combinations of DPF (20×20), TPF (20×20×20) are used. List of BCP features used is: net charge, isoelectric point, secondary structure prediction (DSSP), molecular weight, hydropathy value (kyte-doolitle index), number of hydrogen bond donors and acceptors and the difference between the numbers of hydrogen bond donors and acceptors. Features like frequencies and molecular weight are normalized by the peptide length. To choose the best possible feature vector and to deal with sparse nature of DPF, TPF, we train our model with four sets of features separately and combine the top-scoring features from each set shown in Figure 1. The model combining the top-scoring features is referred to as the Ensemble-Feature (EnF) model. The distributions of lengths, amino acid compositions and biochemical features are shown in SI-S3.

**Figure 1.**
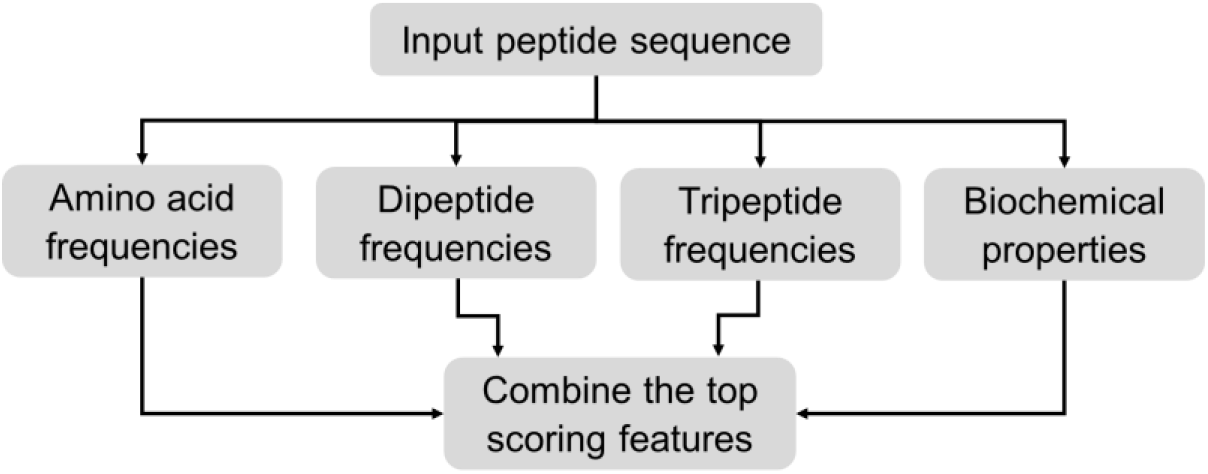
Feature engineering approach adopted in the current work to deal with sparse feature vectors

### 2.4 Metrics used in the current study

To quantitatively measure the performance of the predictors we used following evolution metrics, balanced accuracy, F1 score, sensitivity, specificity, and elements of the confusion matrix, using scikit-learn. Balanced accuracy (Brodersen, K.H. *et al* 2010) is equivalent to normal accuracy with class balanced sample weights. In cases where the classifier does not perform equally well on either class due to an imbalanced test dataset, the balanced accuracy will drop to 1/(number of classes). The evaluation method used was k-fold cross-validation.

### 2.5 Interpreting random forest predictions

For interpreting the random forest predictions, we used the TreeInterpreter package (https://pypi.org/project/treeinterpreter/). This package allows decomposition of each prediction into bias and feature contribution values (Trevor Hastie, Robert Tibshirani, 2009). We analyzed those decision trees which predict ‘CPP’ class from the Test data. We further filter them based on their probability of prediction (at probability >0.65). Decision paths are extracted from these trees. The feature contribution values obtained from these decision paths are plotted against their respective features to study their optimal numbers for predicting the ‘CPP’ class. Due to the random feature selection technique used in RFC algorithms, these values differ across the decision trees, but the overall relative contribution was observed to be consistent in our models.

## 3 Results

### 3.1 Performance of CPP prediction models with different feature vectors

All ML algorithms highly depend on the training dataset and feature vector used in terms of accuracy obtained (Su *et al.*, 2020). Our results with 5 types of feature vectors on 5 types of datasets are shown in Table 1. All the Datasets are divided into Training (80%) and hold-out Test dataset (20%) and the values mentioned in Table 1 are on the Test dataset.

**Table 1.** Performance of CPP prediction models with different sets of features on different datasets. Model AAF - amino acid frequencies, DPF - dipeptide frequencies, TPF-tripeptide frequencies, BCP - biochemical properties, EnF - ensemble featured vector. Dataset A: CellPPD, Dataset B: ‘benchmark’ from CellPPD, Dataset C: SkipCPP-Pred, Dataset D: KELM data, Dataset E: Ensemble data. Balanced accuracy, elements of the confusion matrix and F1 scores obtained on hold-out Test data are mentioned.

| Dataset A | Accuracy (%) | True Positive (%) | True Negative (%) | False Positive (%) | False Negative (%) | F1 score |
| --- | --- | --- | --- | --- | --- | --- |
| Model AAF | 91 | 90 | 92 | 8 | 1 | 0.90 |
| Model DPF | 90 | 88 | 92 | 8 | 12 | 0.89 |
| Model TPF | 86 | 82 | 91 | 9 | 2 | 0.84 |
| Model BCP | 91 | 90 | 92 | 8 | 9 | 0.90 |
| Model EnF | **94** | 92 | 96 | 4 | 8 | 0.95 |
| **Dataset B** |  |  |  |  |  |  |
| Model AAF | 89 | 91 | 90 | 1 | 9 | 0.87 |
| Model DPF | 83 | 83 | 83 | 17 | 17 | 0.77 |
| Model TPF | 83 | 74 | 93 | 7 | 26 | 0.79 |
| Model BCP | 88 | 83 | 93 | 7 | 17 | 0.85 |
| Model EnF | **82** | 74 | 88 | 1 | 12 | 0.75 |
| **Dataset C** |  |  |  |  |  |  |
| Model AAF | 89 | 83 | 95 | 5 | 17 | 0.88 |
| Model DPF | 88 | 82 | 94 | 6 | 18 | 0.87 |
| Model TPF | 83 | 73 | 95 | 2 | 34 | 0.78 |
| Model BCP | 90 | 87 | 94 | 6 | 13 | 0.80 |
| Model EnF | **90** | 85 | 95 | 5 | 15 | 0.89 |
| **Dataset D** |  |  |  |  |  |  |
| Model AAF | 84 | 83 | 86 | 14 | 17 | 0.83 |
| Model DPF | 82 | 75 | 90 | 10 | 25 | 0.80 |
| Model TPF | 76 | 95 | 55 | 45 | 5 | 0.77 |
| Model BCP | 82 | 80 | 84 | 16 | 20 | 0.80 |
| Model EnF | **87** | 86 | 87 | 13 | 14 | 0.86 |
| **Dataset E** |  |  |  |  |  |  |
| Model AAF | 91 | 91 | 93 | 7 | 9 | 0.90 |
| Model DPF | 90 | 88 | 93 | 7 | 12 | 0.89 |
| Model TPF | 87 | 88 | 85 | 15 | 12 | 0.84 |
| Model BCP | 91 | 91 | 92 | 8 | 9 | 0.90 |
| Model EnF | **96** | 92 | 97 | 3 | 9 | 0.93 |

**Table 2.** A brief summary of other predictors.

| CPP prediction model reference | Accuracy | Algorithm | Dataset (number of sequences) |
| --- | --- | --- | --- |
| (A Dobchev, 2010) | 83% | Artificial Neural Network | 100 |
| (Sanders *et al.*, 2011) | 75% | Support Vector Machine | 145 |
| (Gautam *et al.*, 2013) | 94% | Support Vector Machine | 1416<br>(Dataset A in current study) |
| (Chen *et al.*, 2015b) | 83% | Random Forest Classifier | 145 |
| (Tang *et al.*, 2016) | 83% | Support Vector Machine | 925 |
| (Wei, Tang, *et al.*, 2017) | 91% | 2-layered prediction framework based on Random Forest method | 924<br>(Dataset C in current study) |
| (Pandey *et al.*, 2018) | 88 % | Neural Network (Extreme Learning Machine) | 826<br>(Dataset D in current study) |
| Current Prediction model | 96% | Random Forest Method | 2207 |

We get an accuracy of 94.5% for Dataset A which is similar to 92.85% obtained by their (Gautam *et al.*, 2013) support vector machine (SVM) algorithm. We get an accuracy of 90% on Dataset C which is similar to obtained by their RFC and SVM algorithms (Wei, Tang, *et al.*, 2017). With Dataset D, we get an accuracy of 87% which is similar to 86% obtained by their Neural Network algorithm (Pandey *et al.*, 2018). We get an accuracy of 82% on dataset B which was lowest among these datasets.

To understand CPPs at basic amino acid composition level we trained the model with AAF, DPF and TPF. This shows that the average occurrence of positively charged amino acids (Arg, and Lys) is higher in CPPs. While we know this already, we are interested in investigating the contribution of these features in making them cell-penetrating. From our results summarized in Table 1, we obtained highest accuracy with EnF vector followed by AAF/BCP, DPF and TPF, in almost all datasets. The feature vector of DPF and TPF is a sparse matrix as the total number of features are 20×20, 20×20×20 of which most of the values are 0 due to short lengths of peptides. These models do not converge and also have lesser accuracy than others. Hence by taking only important features from them and combining them in EnF, retains neighbouring information and increases accuracy along with helping us understand the preference of selective dipeptides/tripeptides in CPPs.

Five-fold cross-validation accuracy on the training datasets using EnF vector is as following: Dataset A: 0.92 ± 0.03, B: 0.87 ± 0.07, C: 0.89 ± 0.05, D: 0.84 ± 0.08, E: 0.93 ± 0.02. Dataset B and D have lower accuracy values and higher standard deviation. Overall, the performance of the model built with Dataset E using EnF vector was observed to be best in terms of this evaluation method and it also has the lowest standard deviation (more details in SI-S11). We will be using this prediction model for further evaluation and analysis of features. Precision/Recall values calculated on Dataset E are shown in SI-S4.

### 3.2 Testing for the sensitivity of the model towards sequence redundancy and imbalance

To test the sensitivity of the model to sequence similarity, we have built two more prediction models where we remove sequences with more than 80% and 90% sequence identities from the training datasets. We use CDHIT (Huang,Y. et al. (2010) for this reduction. We find that the prediction accuracy decreases by ~10% in these models (SI-S5). CPPs are very sensitive to the changes in amino acid sequences. e.g. 2 mutations in Penetratin makes it a non-CPP (Fischer *et al.*, 2000). If we are too strict with our reduction, we risk losing valuable information and will have smaller data for training Hence our results are not surprising and are actually in agreement with previous studies. (Holton *et al.*, 2013).

To check the sensitivity of our model towards the changes in the size of the classes (CPP and non-CPP), we build prediction models using various resampling techniques. We noticed that these resampling techniques are quite sensitive to the feature vector used and for the current dataset, they do not improve the performance of the model significantly. Results from three resampling techniques (over-sampling the minority class, under-sampling the majority class and SMOTE) are discussed in further details (SI-S6).

### 3.3 Analysis of important features

Figure 2 is a bar graph of feature importance score (FIS) from our EnF model trained on Dataset E (values in SI-S7). The isoelectric point emerges as the property with the highest FIS followed by the ‘difference between the number of hydrogen bond donor atoms and the number of hydrogen bond acceptor atoms’ and net charge. We also find that the top six features i.e. Isoelectric point, the difference between the number of hydrogen bond donor/acceptor atoms, Net charge, Number of Arginines, hydropathy value and molecular weight are sufficient to obtain an accuracy of ~89%. We have not carried out further ‘feature selection’ as the main aim of the study is to understand the contribution of various sequential features in making a peptide cell-penetrating.

**Figure 2.**
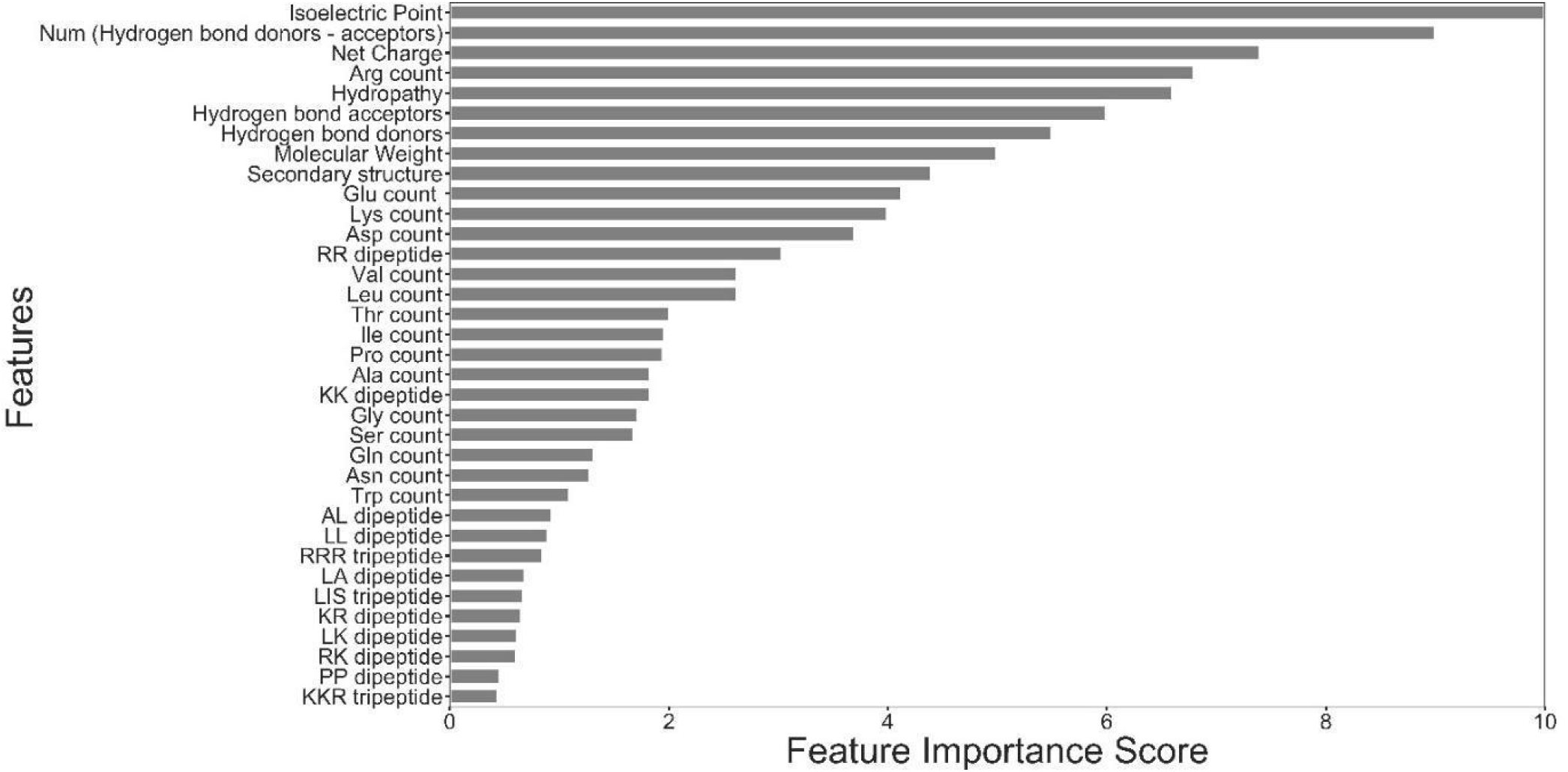
Feature importance score from the EnF prediction model

In our AAF model, amino acids with the highest FIS are Arg, Glu, Lys, Asp, Leu. The 10 most important dipeptides are RR, KK, KR, RK, LA, WK, AL, RW, RI, RL and 10 most important tripeptides are RRR, KKK, KKR, KRK, RRA, RRQ, RKK, RWR, RQR, GRR. This begins to provide guidance for combinations of amino acids to engineer cell-permeability in a peptide; for example, what may be the best amino acid in combination with Arg. Our observations are supported by a few previous studies (Park *et al.*, 2002).

## 4 Discussion

### 4.1 Decision tree path analysis

Interpreting predictions from ML models can be challenging but is an important step to build trust in them and to increase our understanding of the underlying biological phenomena; also observed and discussed in (Yuan *et al.*, 2020). To make our prediction model easily interpretable and intuitive, we used a decision path analysis approach. In RFC, the decision paths from the root of the decision trees to the leaf represent classification rules used by the decision trees to reach a prediction. In order to understand the CPP prediction rules, we need to analyze these decision paths. We have used the TreeInterpreter package as described in Methods for this purpose. The decision tree contribution of each feature is not a single predetermined value but depends on the rest of the feature vector, i.e. all values are relative. Feature vectors determine the decision path that traverses the tree and thus the contributions that are passed along the way. Hence, we obtain a range of contribution values for each feature. Even though these values can vary depending on the members in the Test data, they can guide us in the design of novel CPPs.

### 4.1a Optimal values of biochemical properties

Inferring optimal feature space from ML models can be a difficult task especially for such short peptides and will depend on adequacy of the training data. However, we have attempted to study this using decision path analysis approach. From Figure 3, we observe that isoelectric point above 10, a minimum net charge of 3, hbond donor-acceptor difference of ‘25-50’ and molecular weight of less than 2000 Da have emerged as the optimal range of values to be a CPP. Although it may be argued that several parameters are highly dependent on each other, these numbers do provide good guidance for design purposes.

**Figure 3.**
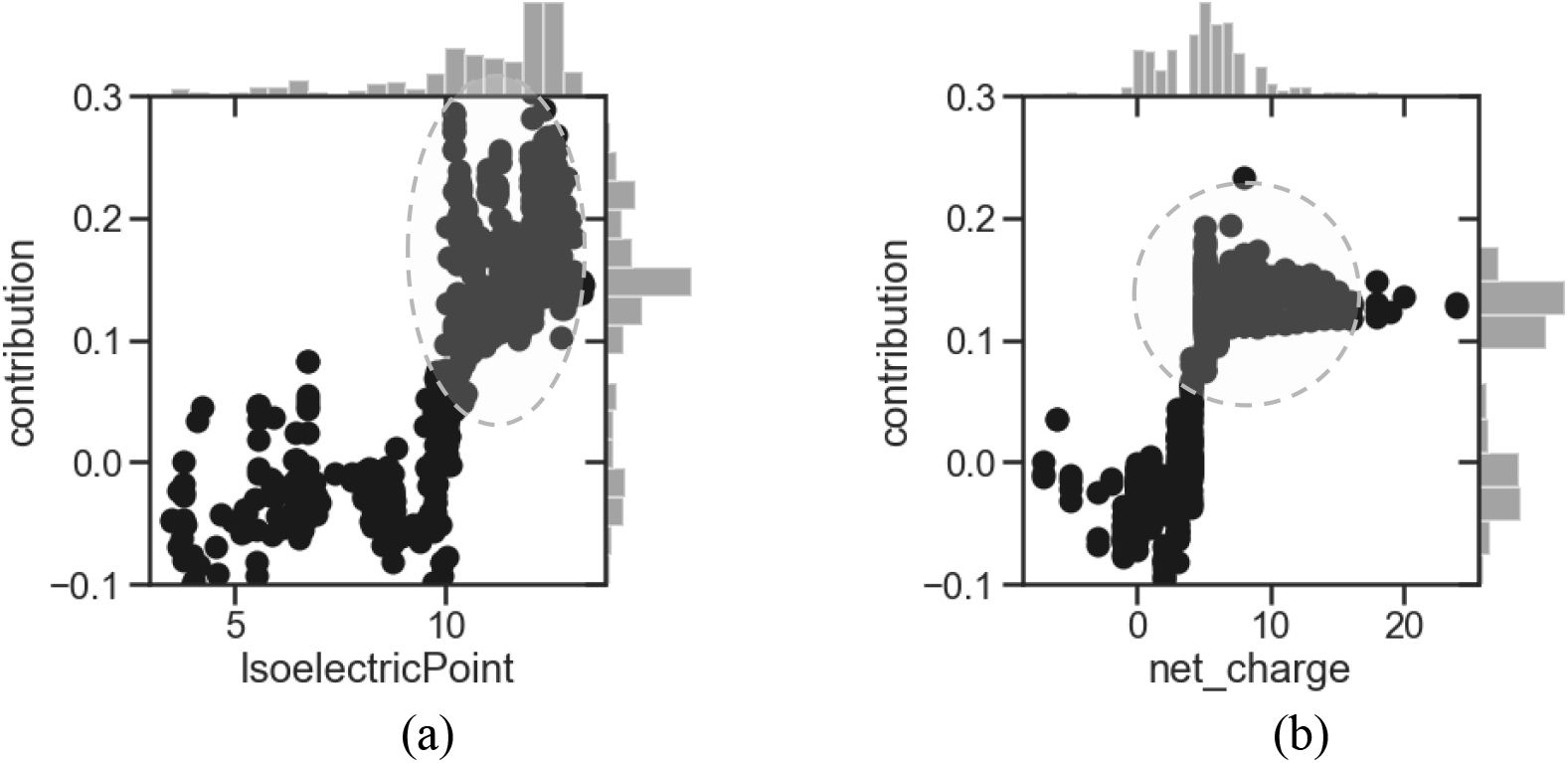
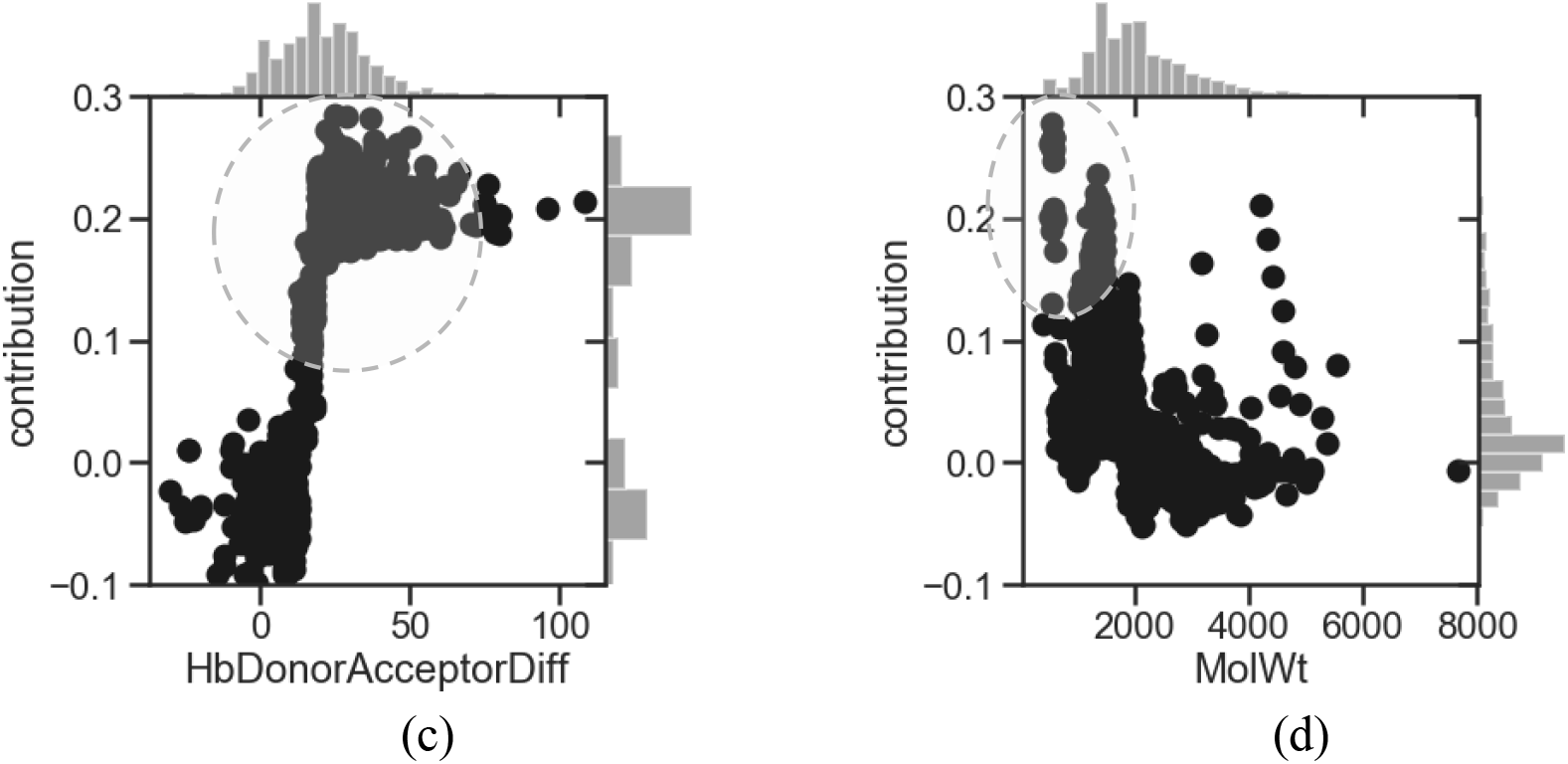
Decision tree path analysis. The contributions obtained from this analysis for important features are shown: (a) isoelectric point, (b) net charge, (c) the difference between the number of hydrogen bond donor and acceptor and (d) molecular weight. X-axis is the value of a feature calculated and each point on Y-axis is the contribution returned by a decision path of a particular decision tree. The proposed optimal feature space is encircled by dotted lines.

### 4.1b Arginines and positive charges in CPPs

The presence of Arg in CPPs has been explored and highlighted for many years and most of the CPPs are Arg rich (Schmidt *et al.*, 2010; Allolio *et al.*, 2018) and as expected, this is been recapitulated in our model too. From our feature contribution analysis (Figure 4) we see that the optimal number of Arg residues for most CPPs is between ~6 and 10 (~17 to 30% of the sequence length). The minimum number of ~6 is also in agreement with previous studies (Wender *et al.*, 2000). In this study Rothbard et. al. prepared a series of TAT peptide (arginine-rich CPP extracted from HIV) mutations and systematically compared their cellular uptake using flow cytometry experiments with those of poly-Arginines of various lengths. The observation that the peptides with high isoelectric points (>10) dominate amongst CPPs (Figure 3) is in accord with the pKa of Arg which is 12.5 (pKa of Lys is 10.5). It has been shown in computational (Yoo and Cui, 2008) as well as experimental studies (Fitch *et al.*, 2015) that Arg predominantly remains protonated under physiological conditions as well as inside a lipid membrane. In addition, the Arg sidechain also engages in the maximum number of hydrogen bonds than other amino acid sidechains and this is likely very relevant for the interactions of the peptide with the phospholipid membranes; for example, the ability to form multiple hydrogen bonds has been shown to be critical for the interactions of Arg-like sidechains with membranes (Fitch *et al.*, 2015; Yoo and Cui, 2008; Li *et al.*, 2013).

**Figure 4.**
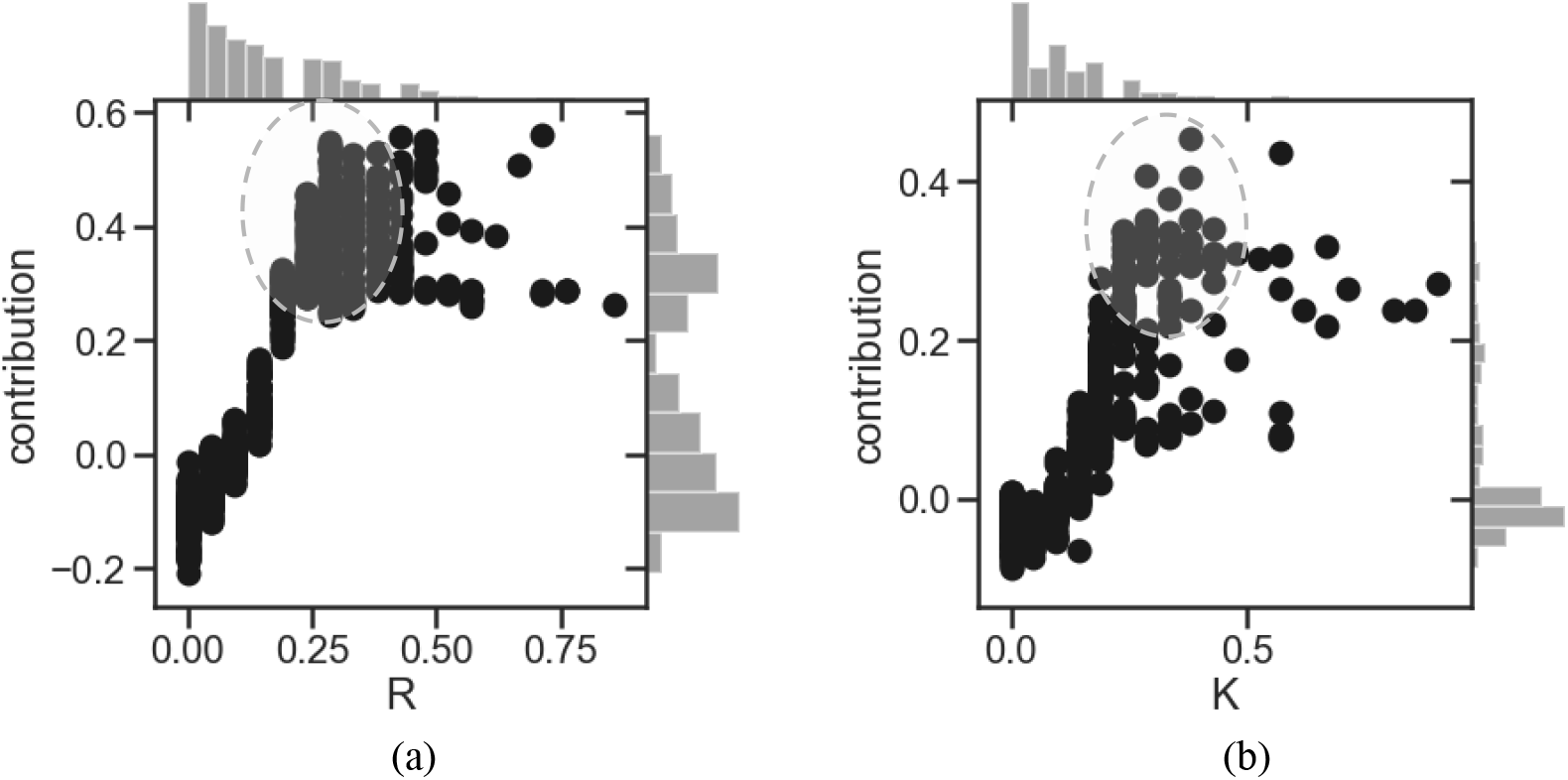
Decision tree path analysis. The contribution values for (a) Arginines (R) and (b) Lysines (K). X-axis is the value of a feature calculated and each point on Y-axis is the contribution returned by a decision path of a particular decision tree. The proposed optimal feature space is encircled by dotted lines.

It was also interesting to observe that Arg, Lys, Leu contribute positively while Glu, Asp contribute negatively (SI-S8). Arg and Glu have similar FIS but opposite feature contribution values. This suggests the absence of Glu might be as important as the presence of Arg in a CPP. The decision path analysis of all other amino acids is discussed in SI-S8.

### 4.1c Prediction model for non-cationic CPPs

Understandably, current CPP datasets are dominant in cationic peptides. So, to understand the characteristics of non-cationic CPPs, we created a dataset of sequences which do not have Arginine and Lysine residues. This dataset has a smaller number of sequences, <200 (the sequences having at least one Arg/Lys are in the range of ~700). These sequences were then divided into 90% for training and 10% for testing and the prediction accuracy obtained was ~90%. Contributions for a few biochemical features extracted from this model are discussed below.

It is clear that in the absence of Arg/Lys, the net charge contributions hint at minimizing the number of negative charges and this is also reflected in lower contributions from the isoelectric point. Interestingly we observe positive contribution from peptides with net-charge near −1 and 0. We also notice that the distribution of molecular weights shifts towards lower values; yet the optimal positive contributions are for molecular weights <2000 Da, as in the case of the full dataset (Figure 3). In the absence of donor atoms from Arg and Lys, the only donors are from Ser/Thr/Asn/Gln/His/Trp sidechains, while the acceptors from Asp/Glu/Ser/Thr dominate and this is reflected in our model. The amino acids with positive feature contributions in this model are Leu, His and Pro (SI-S9). In our model using the full dataset, Leu was observed to contribute positively (following charged residues). Leu is mostly present in amphipathic CPPs such as pVEC. It was also observed that transmembrane helices are enriched in Leu residues and oligoleucines can insert themselves into membranes (Gurezka *et al.*, 1999; Deber and Stone, 2019). It will be interesting to probe further the role of Leu in CPPs. However, we must admit that we have very few non-cationic sequences for this model and the results may change if we manage to find more such sequences.

### 4.2 Comparison with other interpretability methods

We have used two other interpretability methods, LIME (Ribeiro *et al.*, 2016) and SHAP (Lundberg *et al.*, 2018) which are shown to be good methods for explaining ML models. SHAP also uses the TreeInterpreter package as used in this work in their ‘tree explainer’ module. LIME interprets the model locally by fitting a linear model on a locally perturbed dataset while analyzing a decision tree at a time. Contribution values obtained using LIME on one of the datapoints from the Test data is shown in Figure 6 a, these match closely with our analysis. Output obtained using SHAP’s ‘tree explainer’ is shown in Figure 6 b. LIME interprets one decision tree at a time and SHAP provides average feature contribution values whereas our interpretation method provides contribution values for all trees from the RFC model (Figure. 3, 4, 5). The contribution values given by each tree are relative values and depend on the features chosen to build that particular tree, and we may lose information by averaging. We plot values from all decision trees to understand the optimal spaces of each feature; however, we do not capture the interactions between the features. More results from LIME and SHAP are discussed in SI-S10 in further details.

**Figure 5.**
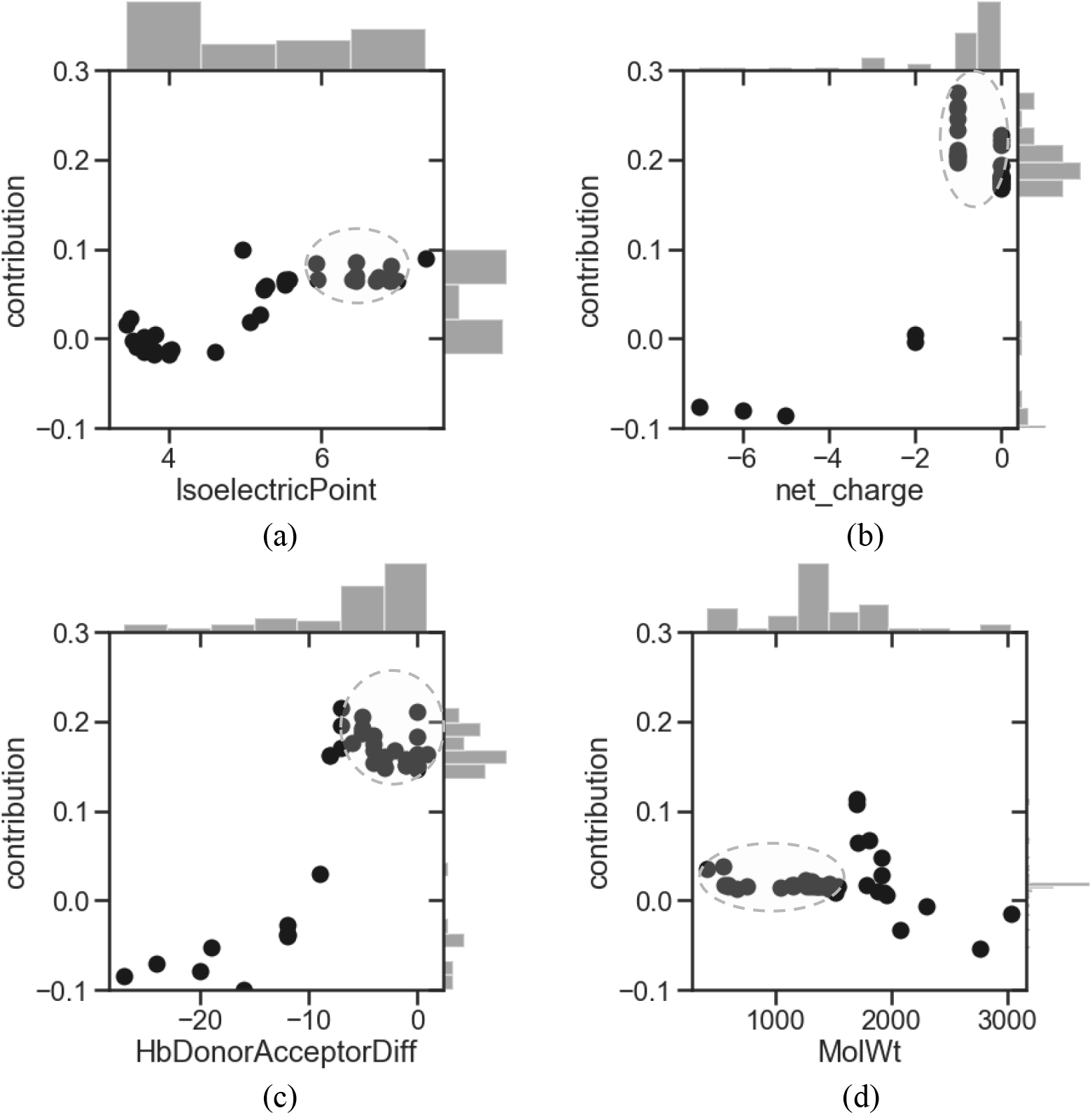
Decision tree path analysis for ‘non-cationic’ CPP prediction model. The contributions obtained from this analysis for important features are shown: (a) isoelectric point, (b) net charge, (c) the difference between the number of hydrogen bond donor and acceptor and (d) molecular weight. X-axis is the value of a feature calculated and each point on Y-axis is the contribution returned by a decision path of a particular decision tree. The proposed optimal feature space is encircled by dotted lines.

**Figure 6.**
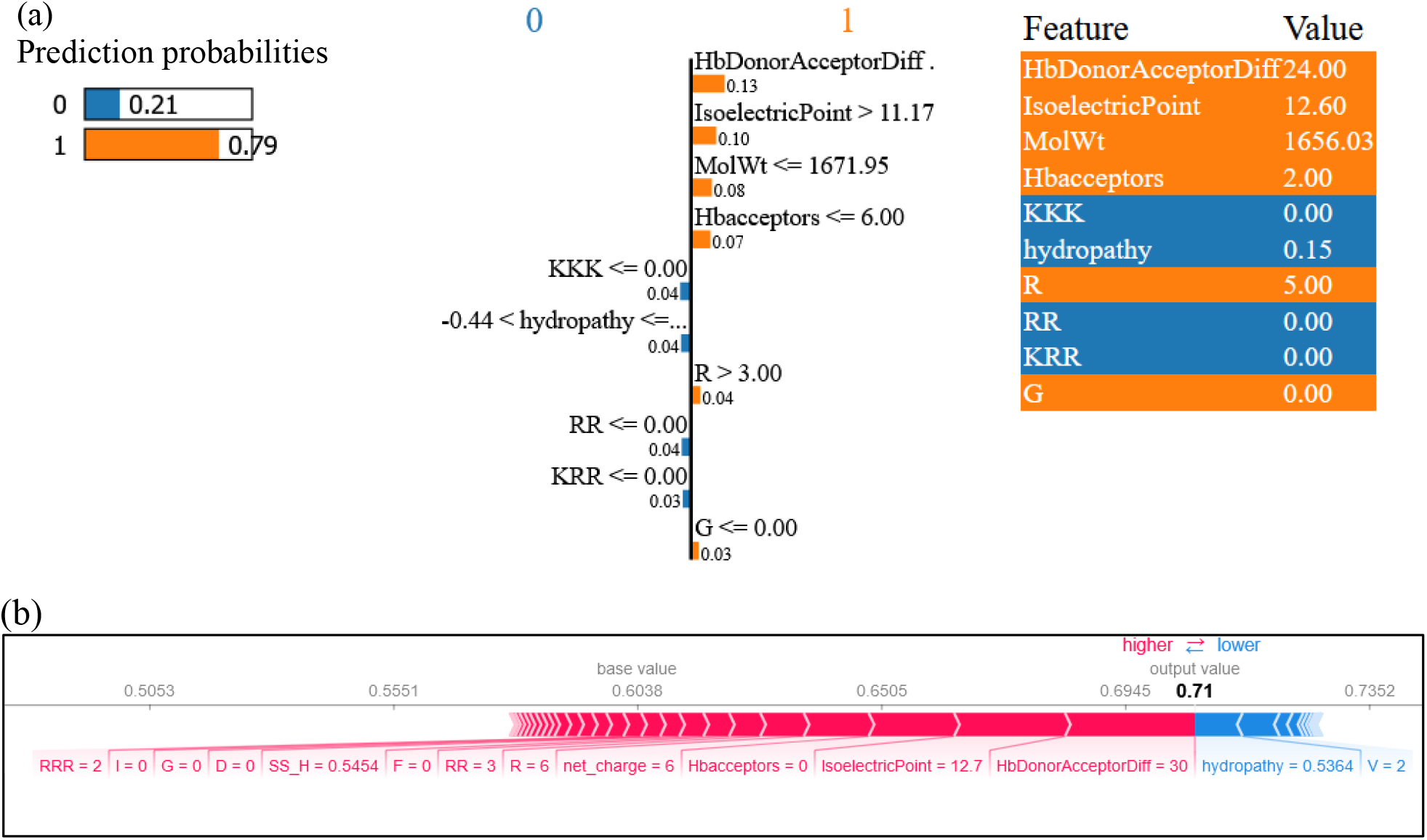
(a) Results from LIME. Column 1 shows the prediction probability for each class (non-CPP - 0 or CPP - 1). Column 2 shows the feature contributions towards each class, with orange colours referring to CPP and blue colours referring to non-CPP. Column 3 lists the properties corresponding to this particular data point. (b) Results from SHAP: Features shown in red have a positive contribution to CPP class when their value is higher. The output value shown in bold is the average prediction probability.

### 4.3 Importance of our Ensemble Feature vector and its role in predicting borderline CPPs

Our EnF model is able to correctly predict peptides which have the same amino acid composition but different sequences and belong to different CPP class. (i) pVEC, a CPP derived from Cadherin, (LLIILRRRIRKQAHAHSK) and scrambled pVEC (Elmquist et al. 2006) (IAARIKLRSRQHIKLRHL) have the same amino acid composition but only pVEC is CPP. (ii) Penetratin vs its non-CPP version (Fischer *et al.*, 2000). These two examples are incorrectly predicted when only AAF or BCP features are used.

### 4.4 Comparison with other prediction models

We observed that combining datasets from different sources resulted in reduced bias and increased quality of the dataset. Table 2 summarizes details of the datasets we have selected. Gautam et al. have used their prediction model to design new CPP sequences, using SVM scores; however, their model is unable to extract specific guidelines in terms of answering questions such as: which amino acids can be incorporated to make a peptide CPP. For this reason, in the current study, we discuss more intuitive design strategies at the basic amino acid property level.

As an additional control, we also build a prediction model with the SVM algorithm of Gautam et al. This SVM model (hyperparameters used after tuning are: RBF kernel, scaled g, c=15, tolerance= 1E-07) applied to their dataset gave us an accuracy of ~96%, which is similar to what they report. The accuracy obtained using SVM on our Dataset E was 90% (with the same hyperparameters mentioned above). It was noted that in the case of SVM, feature scaling/normalization was necessary and without it, the accuracy obtained was only ~75%. One of the main advantages of using RFC is that it requires very minimal data processing and does not require operations such as feature scaling or transformational operations like PCA (Trevor Hastie, Robert Tibshirani, 2009). For this reason, it was easier to rationalize and interpret the feature contribution scores obtained. In addition, even in situations where the dataset is sparse, as is the case with our dipeptide and tripeptide composition matrix, scaling is not recommended (Hoaglin *et al.*, 1983).

### 4.5 Limitations of current prediction models

A major challenge faced by developers of prediction models for CPPs is the quality of available experimental data. These data come from diverse sources and hence are not normalized against the variation in experimental conditions. (i) a high variation in the length of CPPs (5 to 30 amino acids), (ii) a small number of experimentally verified non-CPPs and (iii) variable experimental conditions (concentration, cell lines, etc.) for experimentally validated CPPs; the ability of CPPs to permeate cells depends considerably on these conditions, making the available data hard to integrate into composite datasets. (iv) CPPs enter the cell by various mechanisms and thus have different biochemical properties (Lindgren and Langel, 2011). Some may go inside by passive diffusion; some may go actively (energy-dependent) and hence the properties of the peptide are different for a different class which can add more challenges into prediction. As a result of these issues, the design of a novel CPP is difficult, despite high prediction accuracies in the prediction models. Many sequences in the current training datasets have very similar amino acid compositions. We need a larger variety of CPP sequences and normalized, consistent experimental data. At times the data is mislabeled as it comes from experiments carried out under varying conditions (SI-S2). The need for a large number of validated negative data is also equally important to increase the predictive performance of the algorithms (Wei *et al.*, 2014). Most of the negative sequences in the training datasets come from randomly generated sequences or randomly picked sequences from Uniprot (Gautam *et al.*, 2013; Sanders *et al.*, 2011). Hopefully, the future will witness richer data from experiments on CPPs and on many classes of CPPs, thus improving the training of models and hence their predictive abilities.

## Supporting information

Yadahalli_CPP-SupplementaryInfo

## Acknowledgements

We thank Dr. Roland Huber for carefully reading the manuscript and Dr. Hwee Kuan Lee for discussions on Methods. This work was supported by A*STAR (grant IDs H17/01/a0/010, IAF111213C). SY thanks Varun N.S. for discussions on data processing. We are also grateful to the authors of previous studies who have made their datasets publicly available which enables the current analysis.

## Declaration

CSV is founder Director of Sinopsee Therapeutics and Aplomex; this work has no conflict.

## Notes

### Competing Interest Statement

The authors have declared no competing interest.

