## Supplementary material for "Predicting Cell-Penetrating Peptides: Building and Interpreting Random Forest based prediction Models": Yadahalli_CPP-SupplementaryInfo

**S1. Hyperparameter used in this study for the Random Forest Classifier.**

Hyper-parameters refer to parameters of the estimator algorithm. We carried out this parameter-space tuning using grid search method and optimized it by cross-validation over a parameter grid (using scikit-learn's GridSearchCV). The optimized values are shown below:

bootstrap=True, ccp_alpha=0.0, class_weight= balanced,
criterion='gini', max_depth=30, max_features='auto',
max_leaf_nodes=None, max_samples=None,
min_impurity_decrease=0.0, min_impurity_split=None,
min_samples_leaf=1, min_samples_split=2,
min_weight_fraction_leaf=0.0,
n_estimators=500,
n_jobs=None, oob_score=False

**S2. List of CPP entries with discrepancies in their labels across sources.**

| Sequence | Class | Source |
|---|---|---|
| GRKKRRQPPQC | -1 | Dataset B |
| GRKKRRQPPQC | 1 | Dataset A |
| GRKKRRQPPQC | -1 | Dataset D |
| GWTLNSAGYLLGKFLPLILRKIVTAL | -1 | Dataset B |
| GWTLNSAGYLLGKFLPLILRKIVTAL | 1 | Dataset C |
| GWTLNSAGYLLGKFLPLILRKIVTAL | 1 | Dataset A |
| GWTLNSAGYLLGKFLPLILRKIVTAL | -1 | Dataset D |
| GWTLNSAGYLLGKINLKAPAALAKKIL | -1 | Dataset B |
| GWTLNSAGYLLGKINLKAPAALAKKIL | 1 | Dataset A |
| GWTLNSAGYLLGKINLKAPAALAKKIL | -1 | Dataset D |
| ILRRRIRKQAHAHSK | -1 | Dataset B |
| ILRRRIRKQAHAHSK | 1 | Dataset A |
| ILRRRIRKQAHAHSK | -1 | Dataset D |
| **KIWFQNRRMK** | **1** | **Dataset B** |
| **KIWFQNRRMK** | **-1** | **Dataset B** |
| KIWFQNRRMK | -1 | Dataset D |
| KLALKALKAALKLA | 1 | Dataset A |
| KLALKALKAALKLA | -1 | Dataset D |
| **KLALKALKAALKLA** | **1** | **Dataset B** |
| **KLALKALKAALKLA** | **-1** | **Dataset B** |

| | | |
|---|---|---|
| **KLALKLALKALKAA** | **1** | **Dataset B** |
| **KLALKLALKALKAA** | **-1** | **Dataset B** |
| KLALKLALKALKAA | -1 | Dataset D |
| LLGKINLKALAALAKKIL | -1 | Dataset B |
| LLGKINLKALAALAKKIL | 1 | Dataset A |
| LLGKINLKALAALAKKIL | -1 | Dataset D |
| LLKTTALLKTTALLKTTA | -1 | Dataset B |
| LLKTTALLKTTALLKTTA | 1 | Dataset C |
| LLKTTALLKTTALLKTTA | 1 | Dataset A |
| **LLKTTALLKTTALLKTTA** | **1** | **Dataset D** |
| **LLKTTALLKTTALLKTTA** | **-1** | **Dataset D** |
| LLKTTELLKTTELLKTTE | -1 | Dataset B |
| LLKTTELLKTTELLKTTE | 1 | Dataset C |
| LLKTTELLKTTELLKTTE | 1 | Dataset A |
| **LLKTTELLKTTELLKTTE** | **-1** | **Dataset D** |
| **LLKTTELLKTTELLKTTE** | **1** | **Dataset D** |
| LNSAGYLLGKALAALAKKIL | -1 | Dataset B |
| LNSAGYLLGKALAALAKKIL | 1 | Dataset A |
| LNSAGYLLGKALAALAKKIL | -1 | Dataset D |
| LNSAGYLLGKLKALAALAK | -1 | Dataset B |
| LNSAGYLLGKLKALAALAK | 1 | Dataset A |
| LNSAGYLLGKLKALAALAK | -1 | Dataset D |
| RQIKIFFQNRRMKFKK | -1 | Dataset B |
| RQIKIFFQNRRMKFKK | 1 | Dataset A |
| RQIKIFFQNRRMKFKK | -1 | Dataset D |
| RQIKIWFQNRRM | -1 | Dataset B |
| RQIKIWFQNRRM | -1 | Dataset D |
| RQIKIWFQNRRMKWK | -1 | Dataset B |
| **RQIKIWFQNRRMKWK** | **1** | **Dataset A** |
| **RQIKIWFQNRRMKWK** | **-1** | **Dataset A** |

Class ‘1’corresponds to CPP and Class ‘-1’ corresponds to the non-CPP sequence. Discrepancies within the same source are shown in bold letters. Since these datasets are used in many studies involving CPP prediction or CPP-sequence analysis, we thought it will be useful to show a list of discrepancies which might arrive due to different experimental conditions. (References and description of the datasets is in the main text).

**S3. Distribution of Features used in our Dataset:**

Amino acid frequencies.

<table>
<tr><td>A<br>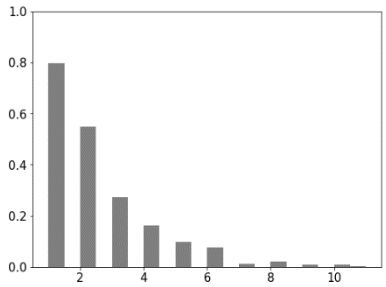
</td><td>C<br>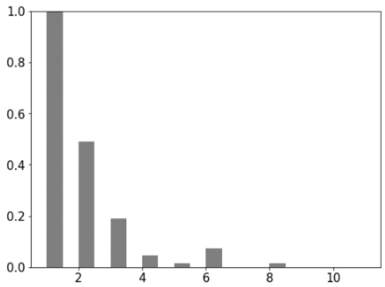
</td><td>D<br>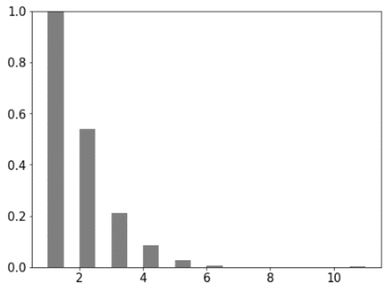
</td></tr>
<tr><td>E<br>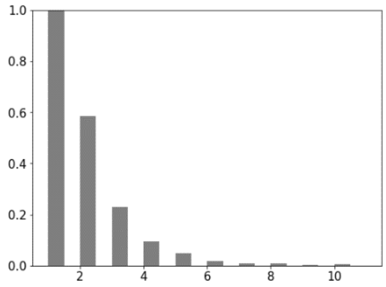
</td><td>F<br>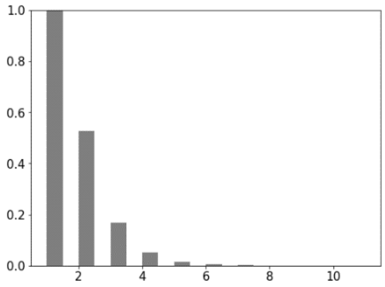
</td><td>G<br>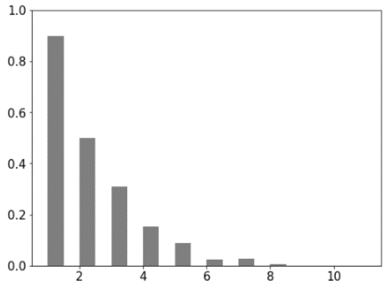
</td></tr>
<tr><td>H<br>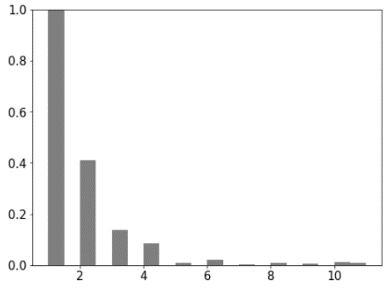
</td><td>I<br>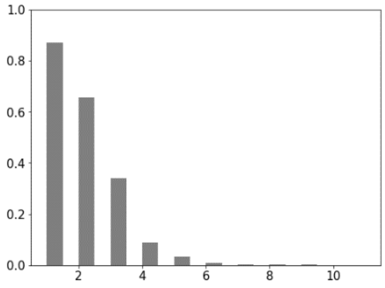
</td><td>K<br>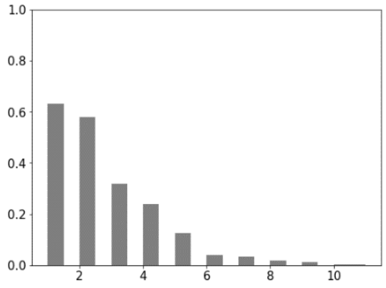
</td></tr>
<tr><td>L</td><td>M</td><td>N</td></tr>
</table>

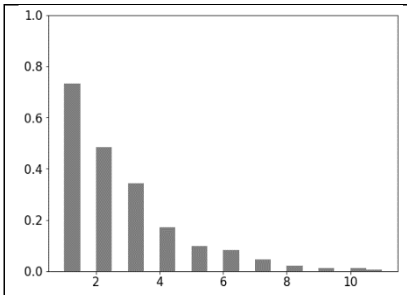
1.0
0.8
0.6
0.4
0.2
0.0
2
4
6
8
10

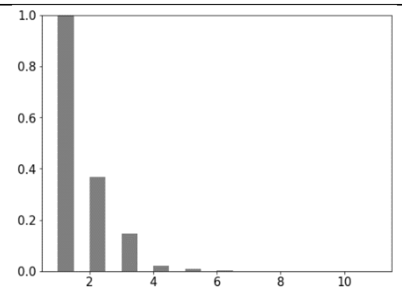
1.0
0.8
0.6
0.4
0.2
0.0
2
4
6
8
10

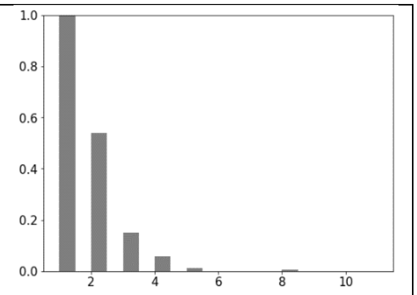
1.0
0.8
0.6
0.4
0.2
0.0
2
4
6
8
10

P

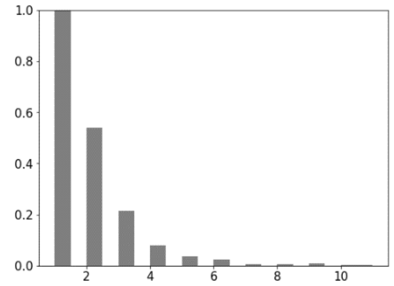
1.0
0.8
0.6
0.4
0.2
0.0
2
4
6
8
10

Q

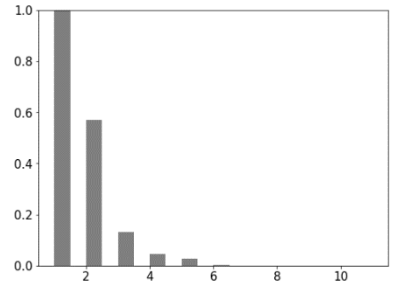
1.0
0.8
0.6
0.4
0.2
0.0
2
4
6
8
10

R

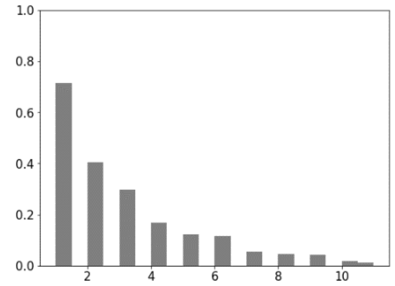
1.0
0.8
0.6
0.4
0.2
0.0
2
4
6
8
10

S

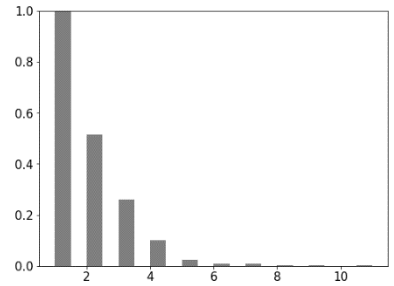
1.0
0.8
0.6
0.4
0.2
0.0
2
4
6
8
10

T

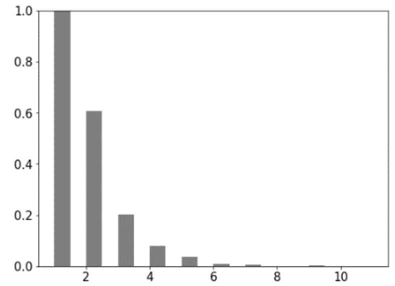
1.0
0.8
0.6
0.4
0.2
0.0
2
4
6
8
10

V

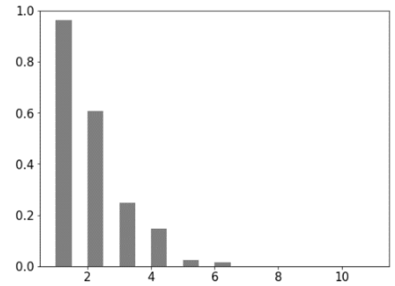
1.0
0.8
0.6
0.4
0.2
0.0
2
4
6
8
10

W

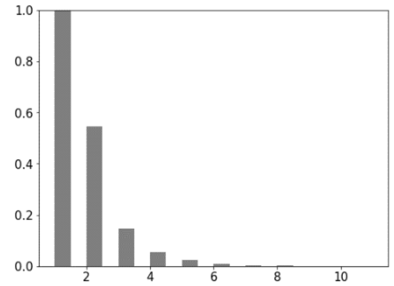
1.0
0.8
0.6
0.4
0.2
0.0
2
4
6
8
10

Y

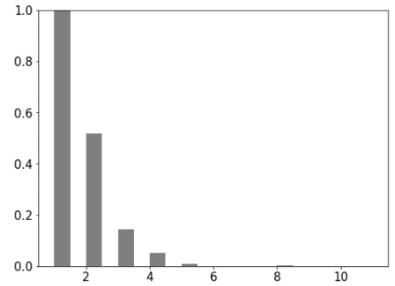
1.0
0.8
0.6
0.4
0.2
0.0
2
4
6
8
10

Biochemical properties

Hydrogen bond acceptors

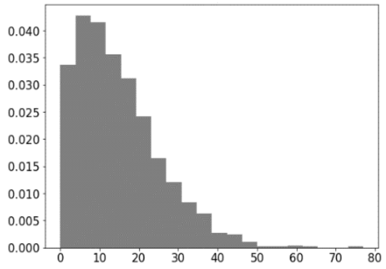

Hydrogen bond donors

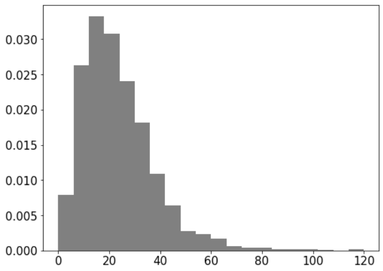

Isoelectric point

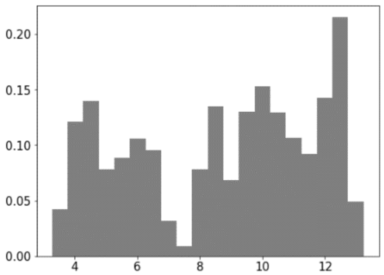

Molecular weight

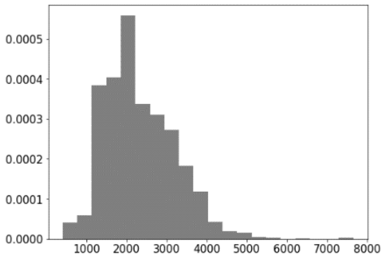

Net charge

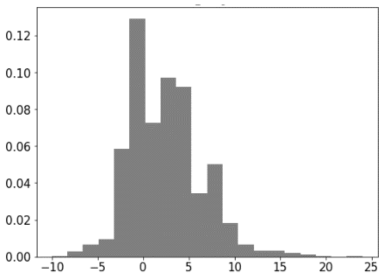

Sequence length

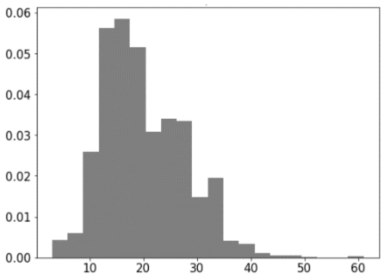

Helical propensity

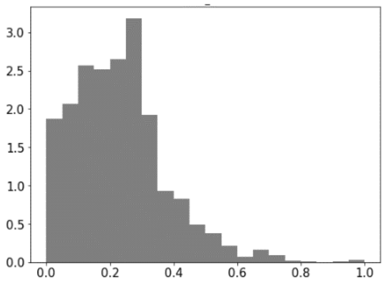

Beta sheet/turn propensity

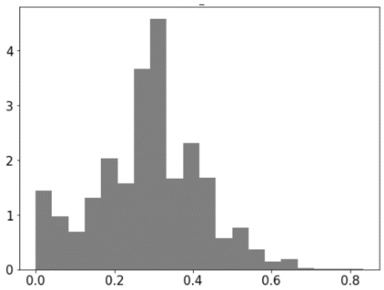

### S4. Classification report.

Classification report for dataset E is shown here. The values are calculated using scikit-learn's 'metrics' module.

| | **Precision** | **Recall** | **F1-score** |
|---|---|---|---|
| Model AAF | | | |
| -1 | 0.93 | 0.93 | 0.93 |
| 1 | 0.90 | 0.91 | 0.9 |
| Model DPF | | | |
| -1 | 0.91 | 0.93 | 0.92 |
| 1 | 0.91 | 0.88 | 0.89 |

| | | | |
|---|---|---|---|
| Model TPF | | | |
| -1 | 0.90 | 0.85 | 0.88 |
| 1 | 0.81 | 0.88 | 0.84 |
| Model BCP | | | |
| -1 | 0.93 | 0.92 | 0.93 |
| 1 | 0.89 | 0.91 | 0.9 |
| Model EnF | | | |
| -1 | 0.91 | 0.97 | 0.94 |
| 1 | 0.96 | 0.9 | 0.93 |

**Results of K-fold Cross-Validation.** Each training process in our prediction models was conducted using five-fold cross-validation. We are showing results of cross-validation on our training data from Dataset E using the Ensemble-feature vector.

Five-fold cross-validation: [0.945, 0.9333, 0.9385, 0.9375, 0.927]

Average accuracy: 0.93 (+/- 0.02)

**S5. Testing for the sensitivity of the model towards sequence redundancy.**

In order to test sensitivity towards sequence similarity in training datasets, we built two more prediction models where we use training datasets after removing sequences with more than 80% and 90% sequence identity. We used CDHIT, a commonly used program, which reduces database sizes by creating a set of representative sequences after clustering them according to a chosen identity cutoff. (Huang,Y. et al. (2010) CD-HIT Suite: A web server for clustering and comparing biological sequences. Bioinformatics, 26, 680–682.)

| Results of the model when sequence identity cutoff of 90% was used | | | | | |
|---|---|---|---|---|---|
| **Dataset E** | **Accuracy** | **True positive** | **True negative** | **False positive** | **False negative** |
| Model AAF | 84 | 71 | 97 | 3 | 29 |
| Model DPF | 84 | 71 | 98 | 2 | 29 |
| Model TPF | 82 | 70 | 95 | 5 | 3 |
| Model BCP | 85 | 76 | 95 | 5 | 24 |
| Model EnF | **86** | 75 | 97 | 3 | 25 |
| Results of the model when sequence identity cutoff of 80% was used | | | | | |
| Model AAF | 82 | 67 | 97 | 3 | 33 |
| Model DPF | 75 | 54 | 97 | 3 | 46 |

| | | | | | |
|---|---|---|---|---|---|
| Model TPF | 74 | 54 | 96 | 4 | 46 |
| Model BCP | 83 | 71 | 97 | 3 | 29 |
| Model EnF | 80 | 62 | 96 | 4 | 38 |

As can be seen from these results, upon the reduction in the size of the database using sequence identity-based redundancy as a criterion, the performance of the model is significantly affected. Accuracy values decrease by more than 10% as compared to the models where the training dataset includes all sequences. Given the functions of these peptides are known to be very sensitive to single amino acid substitutions, this was not very surprising (Holton,T.A. et al. (2013) CPPpred: Prediction of cell penetrating peptides. Bioinformatics, 29, 3094–3096).

**S6. Testing for the sensitivity of the model towards data imbalance.**

The datasets we use are not balanced because the number of non-CPP sequences available is higher than the number of CPP sequences; additional details on these datasets neighbours in the Methods. In order to check if our model is sensitive towards the changes in the number of members from these two classes, we rebuild the prediction models using various resampling techniques as shown below:

1. SMOTE: Synthetic Minority Over-sampling Technique (Chawla, N. V. et al. (2011) SMOTE: Synthetic Minority Over-sampling Technique. J. Artif. Intell. Res., 16, 321–357). New samples are synthesized from the existing dataset by selecting a data point from the minority class at random, find its k-nearest neighbours, and create a synthetic sample near these data points, thus creating new samples near those closer in feature space.
2. Over-sampling the minority class by randomly duplicating a few samples (we also used two resampling strategies, one where all classes are resampled and one where only the minority class is resampled).
3. Under-sampling the majority class by randomly removing a few samples.

We have used scikit-learn's Imbalanced-Learn Library to create these resampling techniques.

| **Dataset E** | **Accuracy** | **True positive** | **True negative** | **False positive** | **False negative** |
|---|---|---|---|---|---|
| **Model AAF** | | | | | |
| SMOTE Oversampling | **92** | 90 | 94 | 6 | 1 |
| OverSampling | **93** | 91 | 95 | 5 | 8 |
| Undersampling | **91** | 90 | 94 | 6 | 1 |
| No resampling (from main text) | *91* | *91* | *93* | *7* | *9* |

| | | | | | |
|---|---|---|---|---|---|
| **Model BCP** | | | | | |
| SMOTE Oversampling | **89** | 88 | 91 | 9 | 12 |
| OverSampling | **91** | 92 | 91 | 9 | 8 |
| Undersampling | **88** | 87 | 90 | 1 | 13 |
| No resampling (from main text) | *91* | *91* | *92* | *8* | *9* |
| **Model EnF** | | | | | |
| SMOTE Oversampling | **90** | 89 | 93 | 7 | 11 |
| OverSampling | **91** | 91 | 93 | 7 | 9 |
| Undersampling | **89** | 89 | 89 | 11 | 11 |
| No resampling (from main text) | ***96*** | *92* | *97* | *3* | *9* |

In the case of feature vector AAF, the model performance is improved by 1-2% but in the case of BCP, the model performance is unchanged. In the case of EnF, accuracy is reduced by 6-7%; it is clear that these re-sampling techniques are quite sensitive to the feature vector used. In summary, for our dataset, the resampling techniques do not improve the performance of the model, this could be because the size of our dataset is not sufficiently large or because our dataset is not highly imbalanced.

**S7. Features in our Ensemble-feature vector and their feature importance scores**:

| | |
|---|---|
| IsoelectricPoint | 0.13372 |
| net_charge | 0.10785 |
| HbDonorAcceptorDiff | 0.10445 |
| Hbdonors | 0.06347 |
| hydropathy | 0.04978 |
| Hbacceptors | 0.04031 |
| R | 0.03982 |
| MolWt | 0.03595 |
| SecStr_Turn | 0.03541 |
| SecStr_Helix | 0.03297 |
| SecStr_Beta | 0.03098 |
| RR | 0.02566 |
| E | 0.02195 |
| K | 0.02133 |
| D | 0.01774 |
| L | 0.01768 |

| | |
|---|---|
| V | 0.0159 |
| T | 0.01477 |
| I | 0.01325 |
| G | 0.01316 |
| A | 0.01312 |
| P | 0.01292 |
| KK | 0.01067 |
| N | 0.01049 |
| S | 0.00916 |
| Q | 0.00831 |
| KKR | 0.008 |
| RRR | 0.0077 |
| LIS | 0.00768 |
| LL | 0.00697 |
| LA | 0.00654 |
| KR | 0.00651 |
| AL | 0.00649 |
| W | 0.00637 |
| KRK | 0.00517 |
| PP | 0.00464 |
| RK | 0.00395 |
| KRR | 0.0038 |
| LK | 0.00371 |
| LPP | 0.00254 |
| WK | 0.00226 |
| KKK | 0.00209 |
| RRQ | 0.0012 |
| LS | 0.00114 |
| WKK | 0.00108 |
| WR | 0.00104 |
| LE | 0.00099 |
| EG | 0.00094 |
| RRA | 0.00093 |
| RWR | 0.00079 |
| RKK | 0.00077 |
| RQR | 0.00075 |
| RIR | 0.00069 |
| EL | 0.00069 |
| RWK | 0.00066 |
| ARR | 0.00064 |

| | |
|---|---|
| EE | 0.00059 |
| VV | 0.00056 |
| DE | 0.00051 |
| WRW | 0.00046 |
| GKK | 0.00037 |

These features are obtained using the feature engineering approach (Fig.1) adopted in the current work.

**S8. Decision tree path analysis on Dataset E** (full data). X-axis is the fraction of amino acid (count of the amino acid/total number of amino acids in a peptide); Y-axis is the contribution value returned by a decision path of a decision tree. The analysis plots for Arg, Lys are shown in the main article. The amino acids with negligible contribution are not shown here.

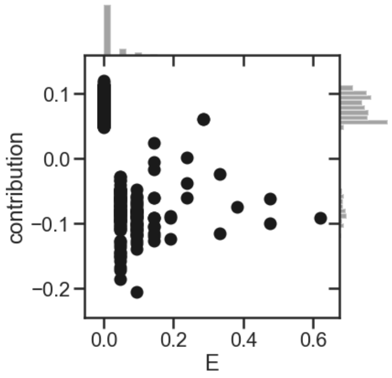

Glu: negatively charged amino acids are not preferred in CPPs; presence of Glu has negative contributions.

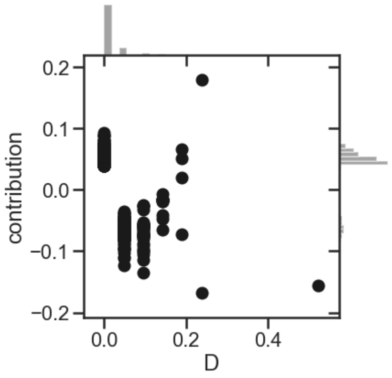

Asp: negatively charged amino acids are not preferred in CPPs; presence of Asp has negative contributions.

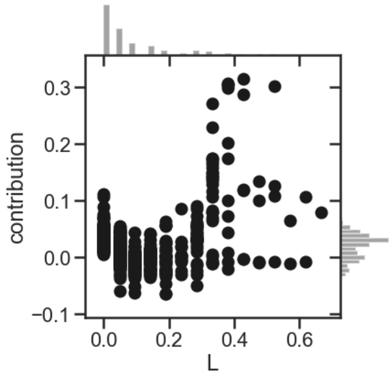

Leu: has positive contributions when fraction is higher than 0.4.

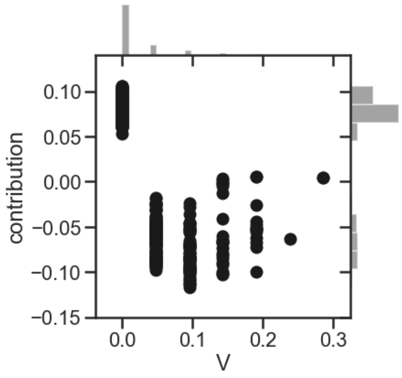

Val: has negative contributions.

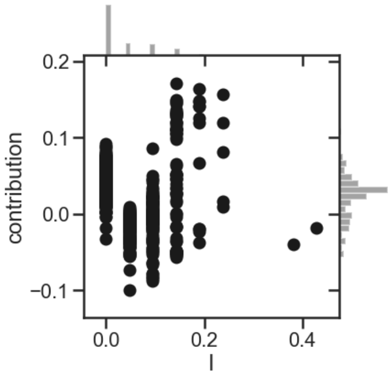

Ile: has neutral contributions for most data points; for fraction higher than 0.1, it contributes positively.

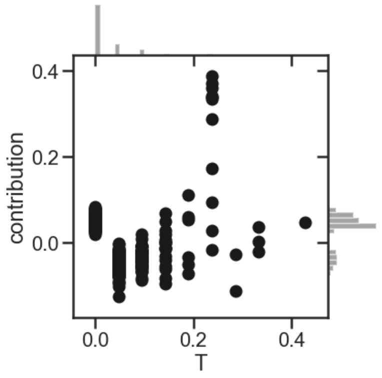

Thr: has neutral contributions for most of the data points; no consistent pattern was found.

| | | |
|---|---|---|
| 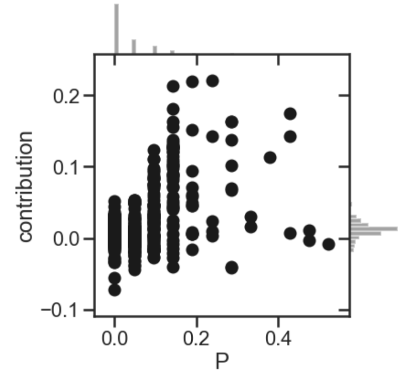  Pro: has positive contributions when fraction is ~0.2, however several outliers were found. | 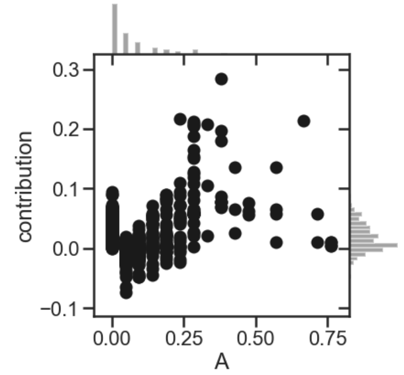  Ala: has positive contributions when fraction is higher than 0.2; optimal value is ~0.3. | 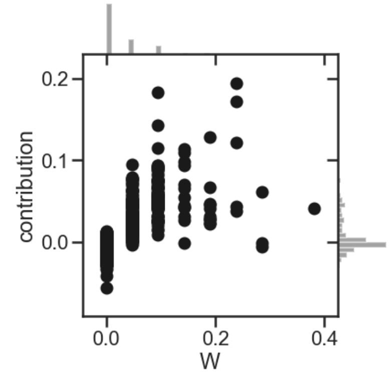  Trp: has positive contributions when fraction is higher than 0.15. |

**S9. Decision tree path analysis of our 'non-cationic' CPP prediction model**. X-axis is the fraction of amino acid (count of the amino acid/total number of amino acids in a peptide); Y-axis is the contribution value returned by a decision path of a decision tree. The amino acids with negligible contribution are not shown here.

| | | |
|---|---|---|
| 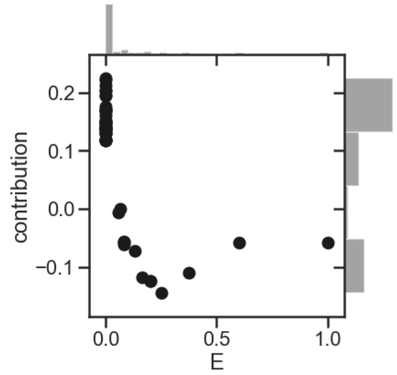  Glu: has negative contributions. | 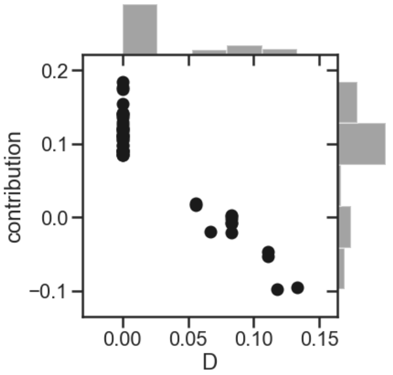  Asp: has negative contributions. | 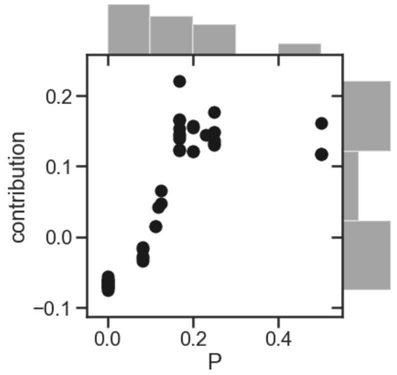  Pro: has positive contributions when fraction is higher than 0.08. |
| 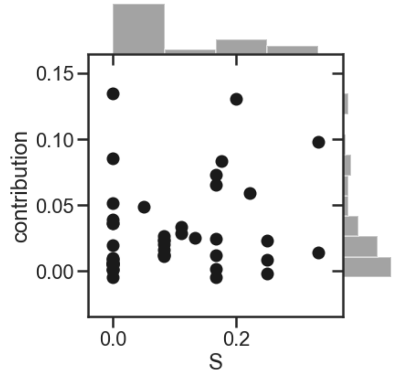  Ser: has positive contributions when fraction is higher than 0.2, but no consistent pattern was found | 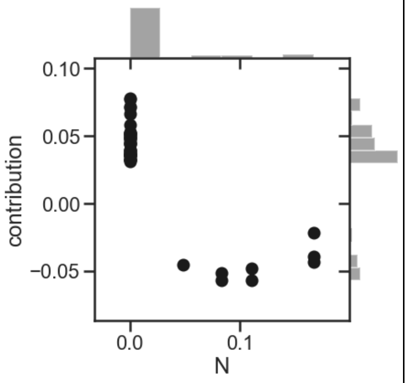  Asn: has negative contributions | 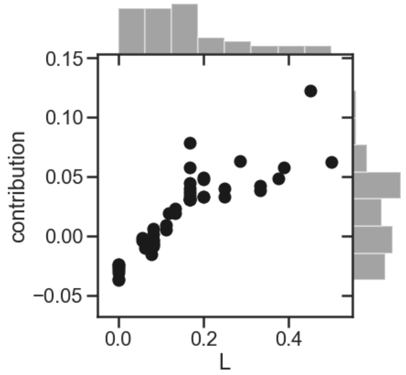  Leu: has positive contributions when fraction is higher than 0.1 |

### S10. Other interpretability methods

#### S10-a. LIME

It is very important to build confidence in our prediction models and to understand why the predictions are right or wrong, particularly in the case of ensemble learning algorithms like RFC. To understand these predictions in an intuitive manner, we have used the ‘Decision tree path analysis’ approach explained in the main text. We have also used two other popular interpretability methods, LIME and SHAP and detail this below.

LIME stands for ‘Local Interpretable Model-agnostic Explanations’ explains the prediction of any classifier in an interpretative manner (Ribeiro T, Marco, Singh S, and Guestrin C. arXiv (2016): "Why Should I Trust You?": Explaining the Predictions of Any Classifier. arXiv-1602.). LIME fits a linear model on a local perturbed dataset while analyzing a single datapoint i.e. a single decision tree in this context. We show below the feature contribution values obtained for amino acid frequencies and biochemical properties upon applying LIME to our models.

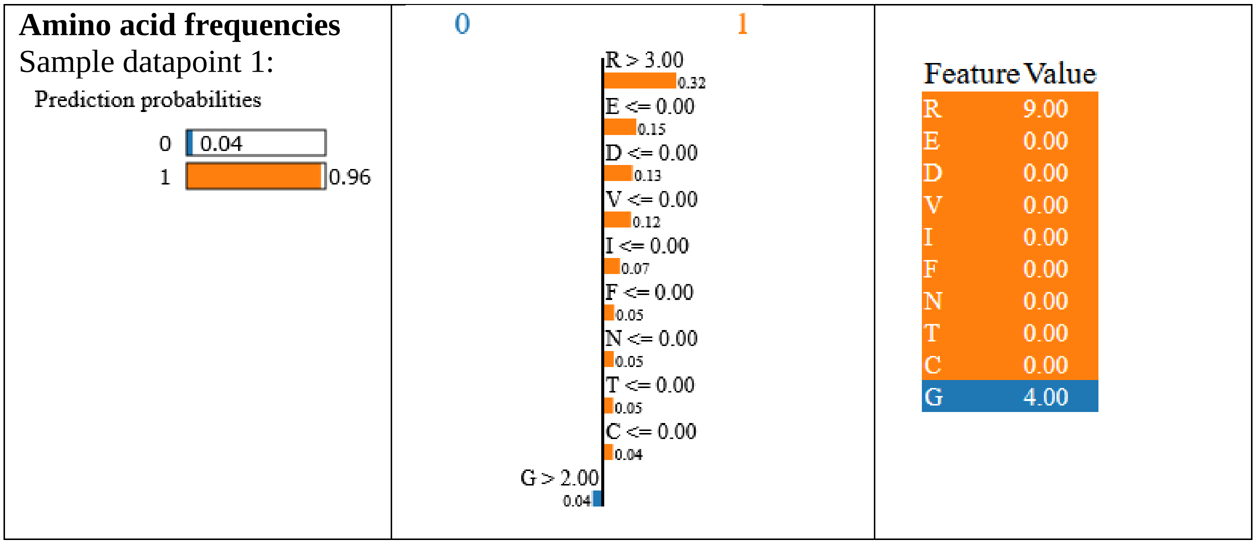

**Amino acid frequencies**
Sample datapoint 2:

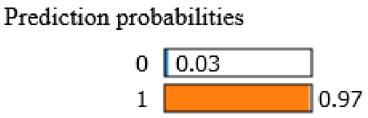

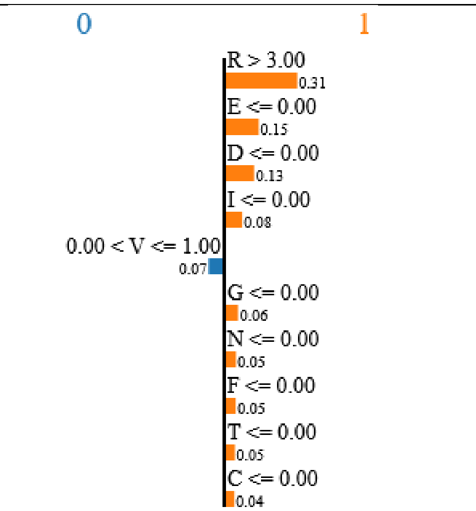

| Feature | Value |
|---|---|
| R | 6.00 |
| E | 0.00 |
| D | 0.00 |
| I | 0.00 |
| V | 1.00 |
| G | 0.00 |
| N | 0.00 |
| F | 0.00 |
| T | 0.00 |
| C | 0.00 |

**Biochemical properties:**
Sample datapoint 1:

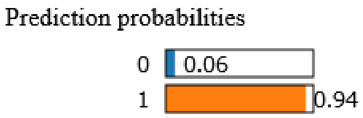

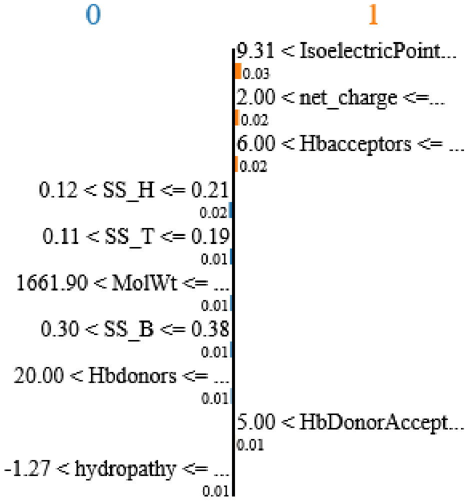

| Feature | Value |
|---|---|
| IsoelectricPoint | 11.00 |
| net_charge | 4.00 |
| Hbacceptors | 8.00 |
| SS_H | 0.20 |
| SS_T | 0.13 |
| MolWt | 1975.28 |
| SS_B | 0.33 |
| Hbdonors | 25.00 |
| HbDonorAcceptorDiff | 17.00 |
| hydropathy | -1.13 |

**Biochemical properties:**
Sample datapoint 2:

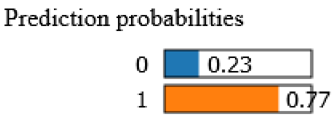

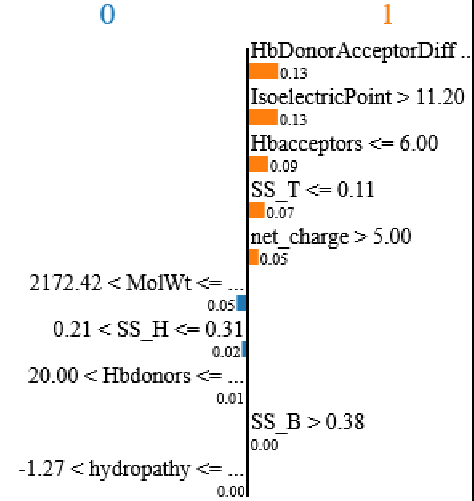

| Feature | Value |
|---|---|
| HbDonorAcceptorDiff | 21.00 |
| IsoelectricPoint | 11.39 |
| Hbacceptors | 4.00 |
| SS_T | 0.11 |
| net_charge | 6.00 |
| MolWt | 2275.84 |
| SS_H | 0.22 |
| Hbdonors | 25.00 |
| SS_B | 0.39 |
| hydropathy | -0.60 |

In this figure, column 1 shows the prediction probabililty for each class (non-CPP - 0 or CPP - 1). Column 2 shows the feature contributions towards each class, with orange colors referring to CPP and blue colours referring to non-CPP (for e.g. Isoelectric point> 11.2, net charge >5, contributes positively to CPP in this particular datapoint; the values shown are feature importance scores. Column 3 lists the properties corresponding to this particular datapoint; orange color refers to positive contributions while blue color refers to negative contribution.

Since it is difficult to understand the analysis of individual data points, we have also used a library from LIME called submodular_pick (SP-LIME). It is a method provided by LIME to select a representative set with explanations; this representative is supposed to provide an intuitive global understanding of the model. Below are the results with SP-LIME, and it has chosen 5 representative samples from our hold-out Test data.

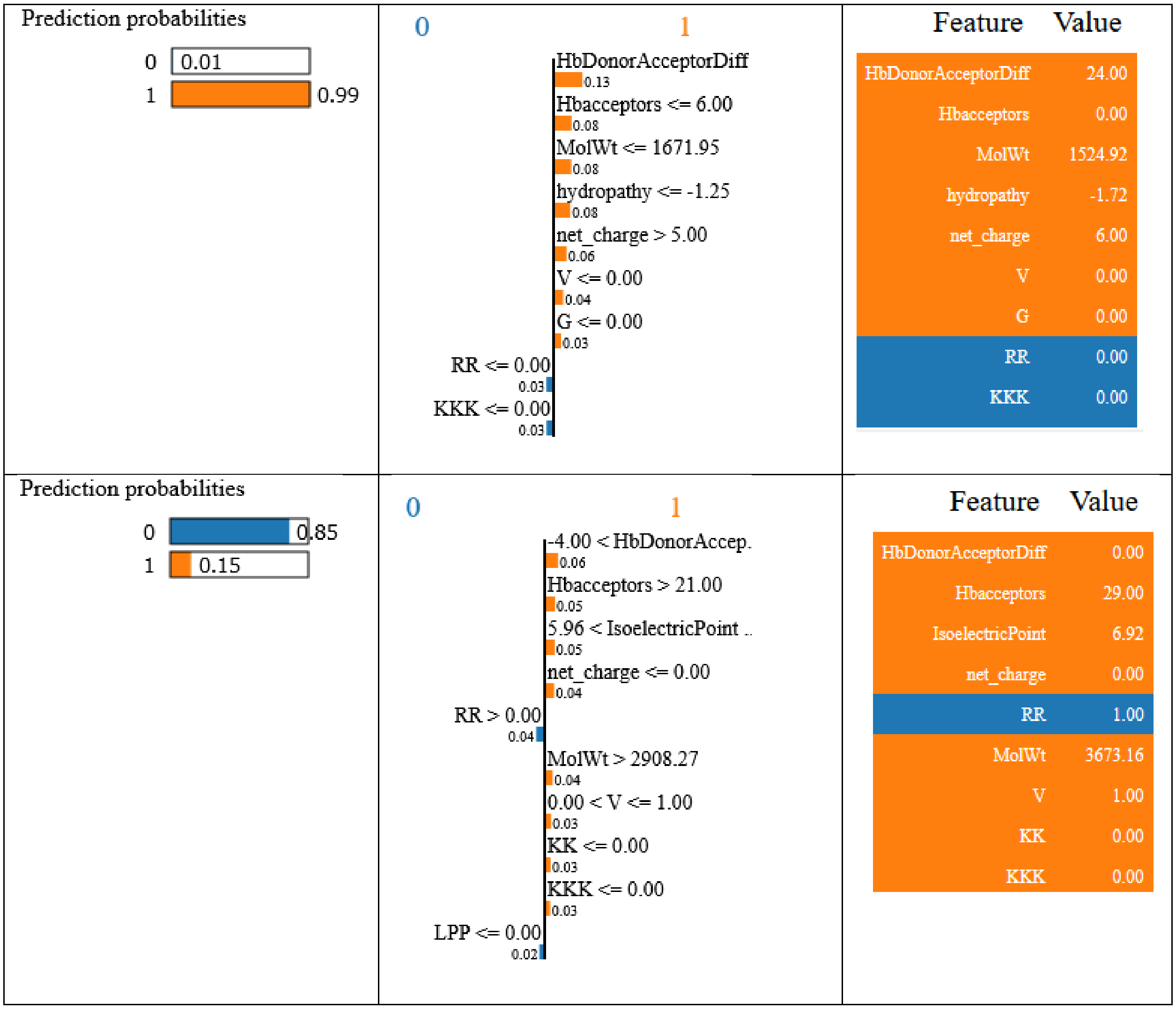

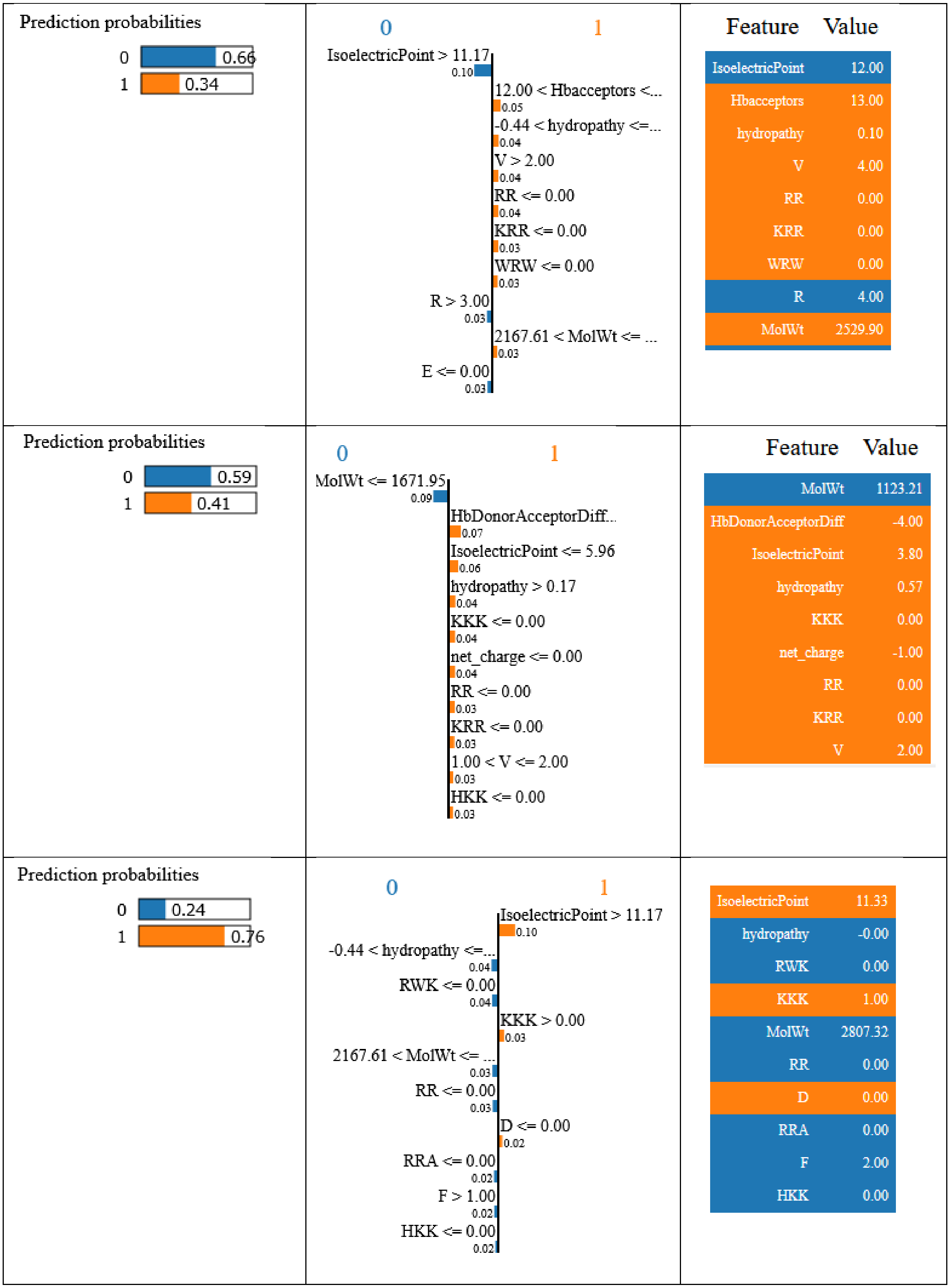

Prediction probabilities
0 0.66
1 0.34
0
1
IsoelectricPoint > 11.17
0.10
12.00 < Hbacceptors <...
0.05
-0.44 < hydropathy <=...
0.04
V > 2.00
0.04
RR <= 0.00
0.04
KRR <= 0.00
0.03
WRW <= 0.00
0.03
R > 3.00
0.03
2167.61 < MolWt <= ...
0.03
E <= 0.00
0.03
Feature Value
IsoelectricPoint 12.00
Hbacceptors 13.00
hydropathy 0.10
V 4.00
RR 0.00
KRR 0.00
WRW 0.00
R 4.00
MolWt 2529.90
Prediction probabilities
0 0.59
1 0.41
0
1
MolWt <= 1671.95
0.09
HbDonorAcceptorDiff...
0.07
IsoelectricPoint <= 5.96
0.06
hydropathy > 0.17
0.04
KKK <= 0.00
0.04
net_charge <= 0.00
0.04
RR <= 0.00
0.03
KRR <= 0.00
0.03
1.00 < V <= 2.00
0.03
HKK <= 0.00
0.03
Feature Value
MolWt 1123.21
HbDonorAcceptorDiff -4.00
IsoelectricPoint 3.80
hydropathy 0.57
KKK 0.00
net_charge -1.00
RR 0.00
KRR 0.00
V 2.00
Prediction probabilities
0 0.24
1 0.76
0
1
IsoelectricPoint > 11.17
0.10
-0.44 < hydropathy <=...
0.04
RWK <= 0.00
0.04
KKK > 0.00
0.03
2167.61 < MolWt <= ...
0.03
RR <= 0.00
0.03
D <= 0.00
0.02
RRA <= 0.00
0.02
F > 1.00
0.02
HKK <= 0.00
0.02
IsoelectricPoint 11.33
hydropathy -0.00
RWK 0.00
KKK 1.00
MolWt 2807.32
RR 0.00
D 0.00
RRA 0.00
F 2.00
HKK 0.00

Our output from the tree interpretation (discussed in the main text) matches closely with the output from SP-L,IME however we look at individual features and their contributions from all decision trees in order to understand the optimal feature space. LIME builds new regression models locally around each data point, so the feature importance and contribution values obtained are different for each point. Hence it is difficult to make global sense of the feature contributions. The R2 score (proportional to confidence value) given by LIME obtained by this local fitting of the model was less than 0.5 in all the cases we have shown here.

**S10-b. SHAP**

We used an additional method, called SHAP, to further interpret our findings. SHAP stands for *SHapley Additive exPlanations*, is a game theoretic approach to explain the output of any machine learning model. The methos is described as "connecting optimal credit allocation with local explanations using the classic Shapley values from game theory and their related extensions" (Lundberg,S.M. et al. (2018) Explainable machine-learning predictions for the prevention of hypoxaemia during surgery. Nat. Biomed. Eng., 2, 749–760). We used SHAP's *tree* explainer on our RFC model and obtained the following interpretation (screenshot provided):

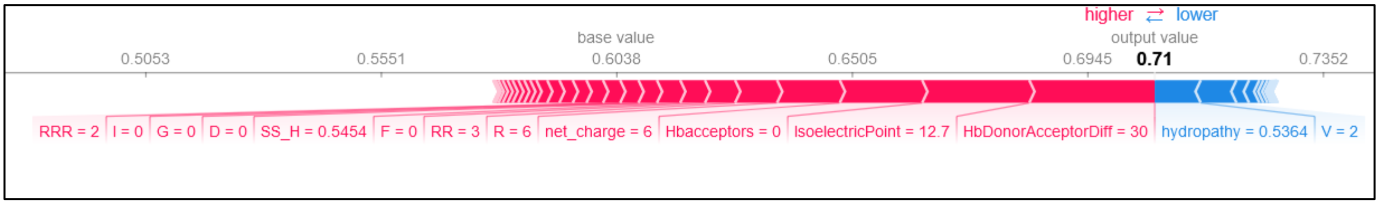

The above figure highlights features which contribute relative to the base values (the mean prediction). 'The value that would be predicted if we did not know any features for the current output, is base value', as defined in their paper. Features shown in red have a positive contribution to CPP-positive class if their value is higher. The output value shown in bold, is the average prediction probability. Overall, SHAP values (labelled on the figures) represent a feature's responsibility for a change in the model output.

In summary, SHAP provides average feature contribution values and LIME interprets one decision tree at a time, whereas our interpretation method provides contribution values for all trees from the RFC model. The contribution values given by each tree are relative values and depend on the features chosen to build that particular tree, and we may lose information by averaging them. Hence, we plot the values from all decision trees to understand the optimal features' space.

### S11. Testing the robustness of Ensemble training dataset.

| Dataset used for training | Accuracy of training dataset | Tested on | Accuracy obtained on test data |
|---|---|---|---|
| Dataset E – Dataset A | 0.88 (+/- 0.04) | Dataset A | 0.91 |
| Dataset E – Dataset B | 0.92 (+/- 0.02) | Dataset B | 0.88 |
| Dataset E – Dataset C | 0.90 (+/- 0.04) | Dataset C | 0.96 |
| Dataset E – Dataset D | 0.91 (+/- 0.02) | Dataset D | 0.85 |
| Dataset E – Dataset B&D | 0.92 (+/- 0.01) | Dataset B/D | 0.84 |

In order to test the robustness of our combined Ensemble dataset (Dataset E), we build four more prediction models. From Dataset E, we omit one dataset (x) at a time and train the model on the remaining sequences i.e. on the sequences from (Dataset E minus Dataset x), and then we use the omitted dataset (x) as a Test. The accuracies obtained on the Test dataset and cross-validation scores on the training dataset are listed the above table. Removal of Dataset B and D results in reducing SD of the Dataset E. This is also in agreement with the results in section 3.1. The accuracy obtained when Dataset B/D are used as Test dataset are also lower. We find that omitting dataset A has the maximum impact on accuracy. In conclusion, it is clear that combining datasets from different sources results in reducing the bias and increased quality of our dataset E.
